# Molecular basis of protein-glycan cross-linking by *Cp*CBM92A revealed by NMR spectroscopy

**DOI:** 10.64898/2026.04.08.717144

**Authors:** Susanne Hansen Troøyen, Marina Susanne Ruoff, Lauren Sara McKee, Gaston Courtade

## Abstract

Our current understanding of carbohydrate-binding module (CBM) function is limited by the fact that most CBM research has focused on single-binding-site modules. CBM family 92 (CBM92) is a recently characterized family of predominantly trivalent proteins that bind β-1,3- and β-1,6-glucans with high specificity. *Cp*CBM92A from *Chitinophaga pinensis* stands out as the first trivalent member of the family to be structurally determined. Multivalent CBM families are rare, and the way in which the three binding sites cooperate in ligand recognition remains unclear. Here, we use NMR spectroscopy to demonstrate how each of the protein’s binding sites plays distinct roles in ligand binding. One binding site, referred to as the β site, can be identified as the primary attachment point because of its higher affinity for all tested ligands, consistent with previous biochemical data suggesting it is the strongest binding site on *Cp*CBM92A. The other two binding sites, referred to as α and γ, preferentially bind longer segments of β-1,3- and β-1,6-glucan chains, respectively. We further show that the glycosidic bond position and anomeric configuration of the binding glucosyl unit strongly affects protein affinity due to a preferred ligand pose in the binding sites. Our results provide insight into how the trivalent architecture of CBM92 might enable cross-linking of scleroglucan chains, which may guide the development of new applications for CBMs in biotechnology.

## Introduction

Carbohydrate-binding modules (CBMs) play an important role in bacterial biomass degradation. They are non-catalytic protein domains typically found in multi-modular enzymes containing one or several carbohydrate-active enzyme domains (CAZymes), where their main function is to aid their enzyme partner in substrate recognition and to maintain enzyme-substrate contact or proximity during catalytic activity [1, 2]. CBMs are classified into families in the CAZy database (www.cazy.org) based primarily on amino acid sequence. The recently established CBM family 92 consists of a group of proteins mostly specific to binding β-1,3 and β-1,6-glucans [3–7], though the first reported CBM92 is a carragenan-binding protein that is now categorized into a small sub-group of the CBM92 family [3, 8]. CBM92 domains are appended to various glycoside hydrolases (GHs) [9] or polysaccharide lyases, most of which are extracellular enzymes secreted by soil and marine-dwelling bacteria to scavenge glycans as carbon sources from biomass [6]. Despite a broad range of substrates targeted by the attached enzymes, which include β-1,3-glucanase (GH16), κ-carrageenase (GH16_17), chitinase (GH18 and GH19), β-1,6-glucanase (GH30), fucosidase (GH95), and α-mannosidase (GH99), most characterized CBM92 modules bind with high specificity to polysaccharides containing β-1,6-linked glucosyl units, like pustulan, yeast cell wall-derived β-1,3-glucan, laminarin, and scleroglucan [7]. Thus, CBM92s appear to largely deviate from the paradigm that CBMs function primarily as substrate binders. An alternative function may involve targeting intact cell walls, where enzyme substrates and glucans with β-1,6-linkages exist in close proximity [10]. Increased enzymatic activity caused by CBMs that target non-substrate glycans has been reported for the pectate lyase Pel10A and xylanases Xyl10A and Xyl10B, where fusion of the enzymes to cellulose-binding CBMs led to increased activity on pectic homogalacturonan and xylan in tobacco stems [11]. Such observations highlight the function of some CBMs in conferring functional advantages beyond direct substrate binding, and these non-substrate-ligand-binding CBMs may be an underexplored resource for biotechnological applications [12].

*Chitinophaga pinensis* is a soil bacterium employing several different CAZymes to break down chitin-rich fungal biomass, which contains considerable amounts of β-1,3 and β-1,6-glucans [5, 13, 14]. One of these enzymes is a GH18 chitinase (UniProt accession number: A0A979GQH9) that is part of a larger multi-modular enzyme containing an N-terminal weakly acting GH5_46 β-1,6-glucanase, and two CBM92 domains — *Cp*CBM92A and *Cp*CBM92B at the C-terminus — that flank the GH18 domain [5]. Li *et al*. demonstrated that the two CBM92 domains do not influence the catalytic activity of the GH18 domain on chitin, but instead contribute to a 10–15°C increased ther-mostability compared to the isolated chitinase [6]. CBMs are well known to enhance the thermostability of CAZymes. This effect has been observed both when comparing native enzymes containing CBMs with their truncated variants lacking these domains, and when CAZymes are fused with CBMs originating from other enzymes [2]. Interestingly, *Cp*CBM92A has been proposed to form cross-links with scleroglucan [7, 15]. This property would make *Cp*CBM92A a highly relevant candidate for biotechnological applications of CBMs, for example in enzyme immobilization and biomaterial design. It is therefore relevant to investigate the ligand-binding properties of this protein to account for structural and biochemical factors that guide polysaccharide interactions.

Both *Cp*CBM92A and *Cp*CBM92B are trivalent CBMs consisting of three distinct subdomains, α, β, and γ (***Figure 1***), with highly conserved amino acid sequences across the CBM92 family [7]. The three subdomains form a β-trefoil structure (***Figure 1***-A), with one glycan binding site in each, all centered around a Trp-Glu motif. They bind specifically to glucans with β-1,6 linkages, including pustulan, laminarin, and scleroglucan. Crystal structures of *Cp*CBM92A, presented by Hao *et al*, showed binding of the non-reducing ends of gentiobiose (Glc-β-1,6-Glc) and sophorose (Glc-β-1,2-Glc) in the three binding sites [7]. The non-reducing end sugar stacks with the conserved Trp residue, and the hydroxyl groups at positions 3 and 4 form hydrogen bonds with the conserved Glu. This binding position could enable extensions from O1 and O6, allowing *endo*-type binding of *Cp*CBM92A along a β-1,6-linked glucan chain like pustulan, or *exo*-type binding to the non-reducing ends of β-1,6-linked oligosaccharides or branches on other polysaccharide substrates.

**Figure 1.**
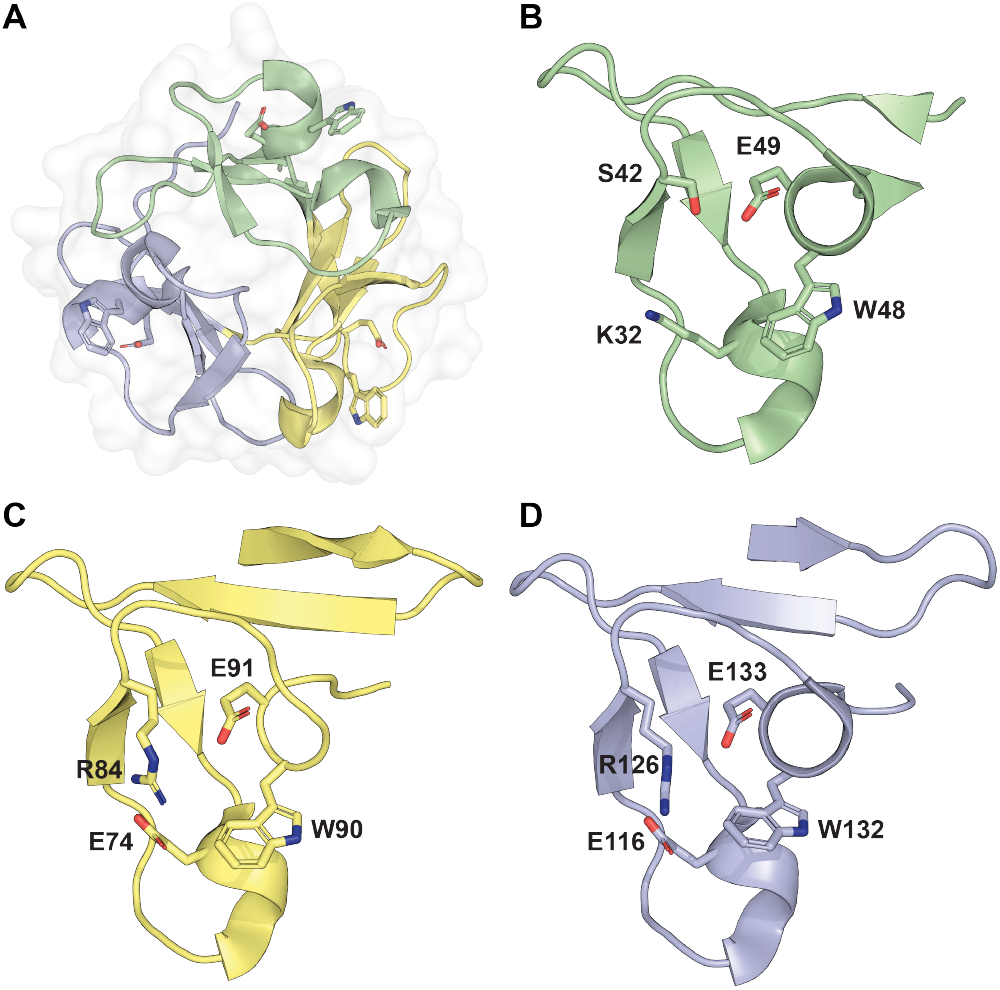
Crystal structure of *Cp*CBM92A (PDB ID: 7ZOI, [7]) (A) and the α (B), β (C) and γ (D) subdomains with key residues in the binding sites shown as sticks.

In this work, we explore the ligand specificity of *Cp*CBM92A in detail using protein- and ligand-observed NMR experiments. Based on the previously established crystal structures, ligand-observed NMR, and Boltz structure predictions [16], we examine the positioning of bound β-1,3- and β-1,6-linked disaccharides and oligosaccharides of glucose in solution, to verify a preferred glucosyl structure for binding to *Cp*CBM92A. We apply protein-observed NMR to simultaneously observe binding at each of the three binding sites without introducing mutations to knock out binding at specific sites, thereby overcoming earlier attempts that were marred by recombinant protein instability and low mutant titers [7]. Our results reveal evidence of different binding modes at the three binding sites of *Cp*CBM92A to oligosaccharide substrates with β-1,3 and β-1,6 linkages. This site-specific variation led us to propose a mechanism for how *Cp*CBM92A might cross-link a branched polysaccharide such as scleroglucan. Deep mechanistic understanding of ligand interaction by multivalent CBMs provides an important addition to our current understanding of CBM function in nature and opens new applications for CBMs in biotechnology.

## Results and discussion

### Chemical shift assignment of the *Cp*CBM92A backbone

We report here the backbone chemical shift assignment of *Cp*CBM92A (HN, H_α_, H_*β*_, N, C’, C_α_, C_*β*_). The ^1^H,^15^N-HSQC spectrum of *Cp*CBM92A with assignments (>84%) is shown in ***Figure S1***. To obtain high sensitivity in triple-resonance spectra, we applied the BEST-TROSY technique [17, 18]. The backbone resonances were assigned based on the HNCA, HNcoCACB and HNCACB experiments. HNCACO, HBHAcoNH and ^15^N-NOESY spectra were used to verify assignments of HN resonances. As the *Cp*CBM92A amino acid sequence contains about 14% Gly, we applied a MUSIC NMR pulse sequence to identify and verify ^1^H,^15^N-HSQC resonances assigned to Gly neighbouring residues [19]. Residues 73–85 belonging to the β subdomain of *Cp*CBM92A could not be assigned to any resonances due to low intensity in the triple-resonance experiments. However, all of the three binding sites Trp48–Glu49 (α), Trp90–Glu91 (β), and Trp132–Glu133 (γ), were successfully assigned. The Trp side chain N_*ε*_H_*ε*_ resonances were identified using a selective MUSIC experiment [20]. Chemical shift data have been deposited at the Biological Magnetic Resonance Bank (BMRB) under the accession number 53615.

### Binding affinities of α, β, and γ sites to glucose and disaccharides of various glycosidic linkages

The potential ability of *Cp*CBM92A to cross-link polymeric chains of scleroglucan is dependent on its trivalency, where each binding site may be involved in forming a network of cross-linked polysaccharide chains. To assess the individual binding affinities of the α, β, and γ binding sites in *Cp*CBM92A to glucose and four β-linked disaccharide ligands, we measured chemical shift perturbations (CSP) in ^1^H,^15^N-HSQC NMR spectra by titrating the ^15^N-labeled protein with each ligand. CSP are a proxy of binding, and consist of changes in the chemical shift caused by alterations in the electronic environment when the protein interacts with a ligand. For most amide groups around the three binding sites, gradual perturbation of the chemical shifts was observed (see ***Figure S2***), indicating fast exchange between the free and bound states [21]. In the β binding site, the ^1^H,^15^N-HSQC signal of Trp90 showed line-broadening instead of CSP, corresponding to slower exchange and higher affinity resulting from a lower off-rate *k*_off_. Dissociation constants *K*_D_ (***Table 1***) were estimated for each site by averaging *K*_D_ values for central residues with high CSP in each binding site. The β binding site had the highest affinity of all sites. This aligns with earlier site-directed mutagenesis experiments removing the primary binding Trp residues from each subdomain in *Cp*CBM92A, which revealed that the β binding site is the most important for binding to β-1,3-glucan, scleroglucan, and laminarin [7]. In the current work, the β and γ sites are determined to have about 2-fold higher affinity to gentiobiose (Glc-β-1,6-Glc) than to the other disaccharides (discussed below). The α site displayed 2-fold higher affinity to cellobiose (Glc-β-1,4-Glc) than to the other disaccharides. This indicates that the alpha site may be more promiscuous, due to the less bulky Ser42 shaping the binding site, compared to the arginine residues (Arg84 and Arg126) present in the β and γ subdomains (***Figure 1***). These arginine residues have been proposed by Hao *et al*. to contribute to higher binding strength, as knock-out mutants depleted of the β and γ Trp residues showed a more pronounced decrease in binding ability than the α knock-out variant [7]. Our results show that, despite having some preference for the type of glycosidic linkage, all binding sites of *Cp*CBM92A are able to bind all four β-linked disaccharides of glucose, with the β-binding site in particular showing the highest affinity for the Glc-β-1,6-Glc linkage found in the protein’s preferred polysaccharide ligands.

**Table 1.** Dissociation constants *K*_D_ for the interaction of *Cp*CBM92A α, β and γ binding sites with glucose and disaccharide ligands, determined from CSP in ^1^H,^15^N-HSQC NMR spectra.

| Ligand | $K_d$, $\alpha$ (mM) | $K_d$, $\beta$ (mM) | $K_d$, $\gamma$ (mM) |
| --- | --- | --- | --- |
| Glucose | $37 \pm 2$ | $1.8 \pm 0.1$ | $14 \pm 1$ |
| Sophorose (β-1,2) | $7.1 \pm 0.6$ | $1.5 \pm 0.1$ | $9.6 \pm 0.4$ |
| Laminaribiose (β-1,3) | $8.1 \pm 0.5$ | $1.78 \pm 0.01$ | $7.0 \pm 0.6$ |
| Cellobiose (β-1,4) | $3.6 \pm 0.2$ | $1.36 \pm 0.07$ | $11.3 \pm 0.1$ |
| Gentiobiose (β-1,6) | $9.2 \pm 0.4$ | $0.68 \pm 0.03$ | $4.6 \pm 0.1$ |

### WaterLOGSY NMR experiments reveal a preferred local ligand structure at the binding sites

Crystal structures of *Cp*CBM92A presented by Hao *et al*. showed binding of the non-reducing end of gentiobiose and sophorose in the binding clefts, with an orientation that could allow extensions on O1 and O6 of the binding sugar [7]. To probe the sugar structure selectivity of *Cp*CBM92A and local ligand positioning at the Trp binding site, we investigated the interaction of glucose, laminaribiose (Glc-β-1,3-Glc), and gentiobiose with *Cp*CBM92A in solution using WaterLOGSY NMR. This ligand-observed NMR experiment enables the determination of the solvent accessibility of each proton in the ligand upon binding to the protein. Protons from a fast-tumbling free ligand give rise to negative signals in the WaterLOGSY spectrum, as a result of NOE transfer from bulk water. However, when the ligand is bound to a protein, NOE transfer from water molecules on the protein surface results in more positive signals. Usually, the bound ligands will have an additional positive contribution from indirectly relayed spin diffusion via the protein, but we did not detect this effect in our experiments, as *Cp*CBM92A is too small to have a high contribution from spin diffusion. The solvent-accessibility of the ligands was quantified through the WaterLOGSY factor, WF, which is the relative change in signal intensity in the presence and absence of protein [22]. Ligand proton chemical shifts were assigned based on the observed *J*-coupling constants and comparison with previously reported chemical shifts [23] (see ***Figure S3, Figure S4, Figure S5*** and ***Table S1***). We observed high WF at the non-reducing end of both disaccharides, corresponding to lower solvent accessibility of these protons (***Figure 2***). To supplement the WaterLOGSY experiments, we also measured ligand-observed CSP by comparing chemical shifts in the absence and presence of protein. CSP results not only indicated the same binding effect at the non-reducing end (***Figure S6***) but also confirmed an equally strong binding effect on the reducing end of β-gentiobiose, which also showed high WF. The molecular structures of the reducing and non-reducing ends of β-gentiobiose are similar (***Figure 2***-B); they both consist of six-membered rings with extensions on position 5 and/or 1 adjacent to the oxygen atom of the ring, and they both have β configuration at the anomeric carbon. This structural similarity further suggests that this specific ligand structure may be essential for binding to the Trp-Glu residues at all three binding sites. In addition, it explains the 2-fold higher affinity of the β and γ sites for gentiobiose compared with the other tested disaccharides, in which only one of the two glycosyl units harbors this preferred structure. As glucosyl units with α configuration at the anomeric carbon displayed lower WF and CSP for gentiobiose and glucose (***Figure S7*** and ***Figure S8***), we conclude that there is a strong preference for binding β-anomers. This is likely due to a weakened stacking interaction of α-anomers with the binding Trp residue imposed by the axial OH group, and aligns well with previously published data showing no binding of *Cp*CBM92A to starch [7].

**Figure 2.**
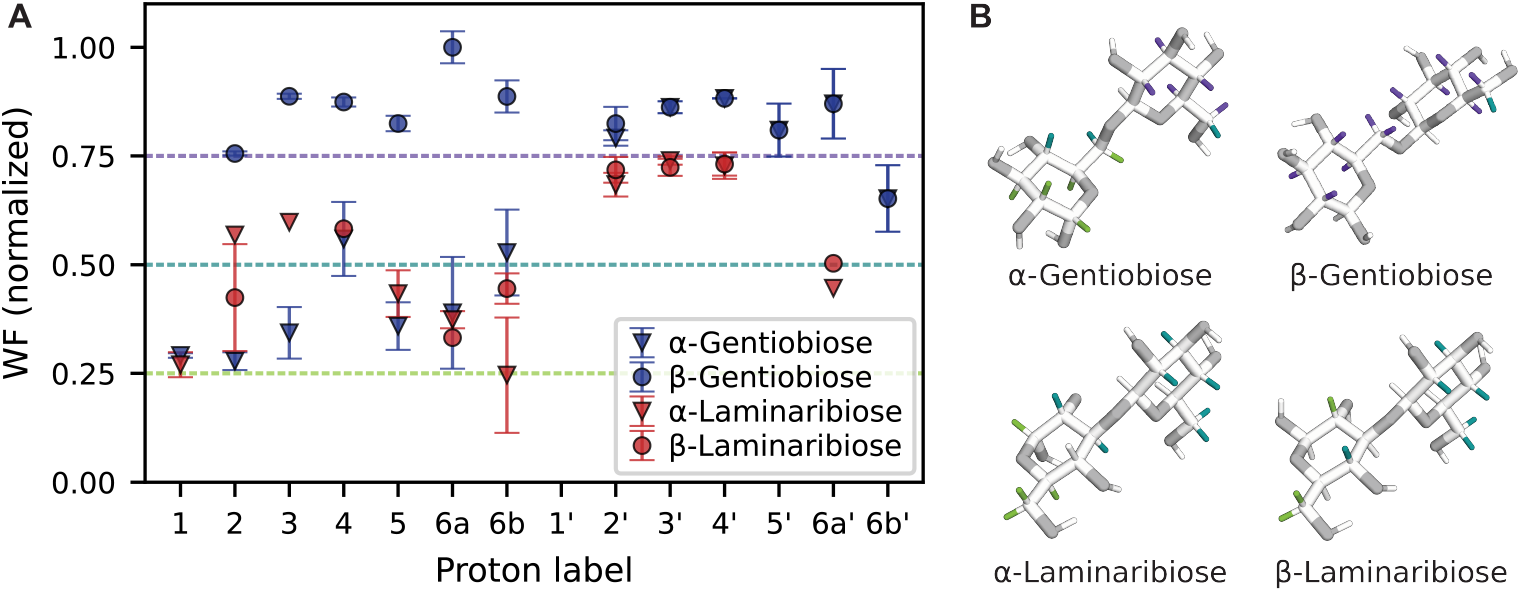
WaterLOGSY factors (WF) of protons of laminaribiose and gentiobiose binding to *Cp*CBM92A (A). Protons of the non-reducing end are marked with ‘. All WF values were normalized to the maximum WF. Error bars represent the standard deviation of WFs calculated for each line picked for a specific proton (see ***Figure S9*** and ***Figure S10***). Ligand structures (B) with protons colored according to WF <0.25 (light green), <0.50 (teal), and <0.75 (dark purple).

### Prediction of protein-ligand complex structures

We used the deep learning model Boltz-1x to predict the structures of *Cp*CBM92A in complex with glucose and disaccharide ligands [16, 24]. For most of our ligands, the model incorrectly interpreted stereochemistry, an issue previously reported with Boltz-1 and AlphaFold3 [25, 26]. To circumvent this, we used Gnina to re-dock the correct ligand structures in the binding sites identified by Boltz-1x, and to predict their affinities [27, 28]. Although the α, β, and γ binding sites were correctly identified in Boltz-1x, the ligand orientation varied in separate runs of Gnina (***Figure S11***). Still, the dockings reflected some of the observations from our NMR experiments. Disaccharides docked in the α binding site more frequently showed extensions in the space around Ser42 (see ***Figure 1***), indicating that this site may be more promiscuous with regard to ligand orientation. Gnina applies a convolutional neural network (CNN) scoring function to rank poses of docked ligands based on the plausibility of their 3D structure [28]. Higher CNN scores were consistently observed for the β binding site than for the other two sites (***Figure S12***), which possibly reflects the importance of this binding site and its favorable topology for binding small ligands. However, the *K*_D_ predicted by Gnina (CNN affinity) did not distinguish the β site as a higher-affinity binding site (***Figure S13***). We performed 20 iterations of Gnina dockings per ligand, and binding at the non-reducing end was observed in 66% of the top 3 scoring poses (***Figure S12***). Our WaterLOGSY experiments showed that the reducing end of β-gentiobiose was able to bind *Cp*CBM92A (***Figure 2***), and in the top-scoring (CNN scores) structures of β-disaccharides, only β-gentiobiose docked at the reducing end (at the β binding site). Finally, the docking poses illustrated the two possible positions of the extensions from the binding glucosyl unit at O1 and O6, as shown in ***Figure 3***. The predicted orientation of the binding glucosyl unit of laminaribiose and gentiobiose in the β binding site matches the previously reported crystal structure [7].

**Figure 3.**
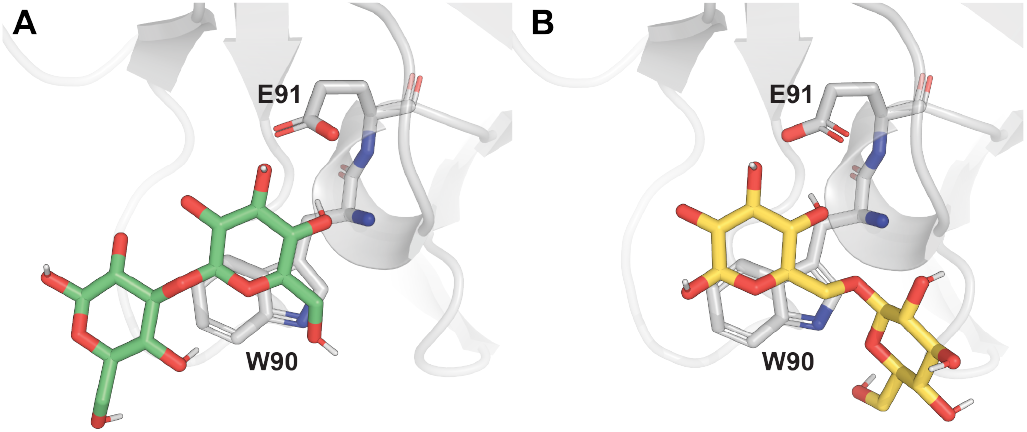
Predicted structure of laminaribiose (A) and gentiobiose (B) docked to β binding site of *Cp*CBM92A, showing the directions of possible extensions from the binding glycosyl unit on O1 and O6. The protein-ligand complex structure was predicted using Boltz-1x, and the ligands were re-docked to the binding site using Gnina, to obtain the correct stereochemistry.

### *Cp*CBM92A binds *exo* to laminarihexaose and *endo*/*exo* to gentiohexaose

Based on the disaccharide experiments and structure predictions suggesting a preferred structure of the glycosyl unit interacting with the binding site, we probed the interaction of *Cp*CBM92A to longer oligosaccharide ligands with β-1,3 and β-1,6 linkages; laminarihexaose and gentiohexaose. For easier referencing, the glycosyl units of the hexamers are labeled with the letters A-F, with A representing the reducing ends, and F the non-reducing ends (see ***Figure S14***). Proton chemical shifts and assignments are shown in ***Table S2***. Chemical shift assignment of the anomeric protons of laminarihexaose was based on [29]. To observe all the ligand resonances in the anomeric region of the NMR spectra (around 5.3-4.5 ppm), these experiments were conducted at lower temperatures (8.9 °C). The lower temperature, possibly combined with a higher ligand affinity, shifted the exchange kinetics from the fast exchange regime observed for the disaccharides into an intermediate exchange regime. Consequently, the dominating binding effect observed in the ^1^H spectra was increased line-broadening in the presence of *Cp*CBM92A (***Figure 4***). This effect arises from increased transverse relaxation rates *R*_2_ of the affected nuclei due to binding to a large macro-molecule with a slower tumbling rate in solution, and chemical exchange between the free and bound states. For laminarihexaose (***Figure 4***-A), very strong line-broadening was observed for resonances at the non-reducing end (F), shown by almost undetectable, broad signals at 4.77 ppm (H1-F), 3.37 ppm (H2-F), and 3.41 ppm (H4-F), which were all well-resolved in the spectrum. No binding effect in the form of line broadening or CSP could be detected at the reducing end (A) or the adjacent glycosyl unit (B), which both showed sharp doublets for the anomeric protons at 5.24 ppm (H1-A, α), 4.68 ppm (H1-A, β), 4.79 ppm (H1-B, α) and 4.76 (H1-B, β), and a sharp triplet at 3.44 ppm (H2-A) in the presence of the protein. This means that *Cp*CBM92A has a strong preference for *exo* binding at the non-reducing end, which is the only glycosyl unit in the oligomer with no extension on position 3. For gentiohexaose (***Figure 4***-B), strong line-broadening was observed for all resonances, showing that all glycosyl units of this ligand are involved in binding, including the non-reducing end. The observation of both *endo* and *exo* binding to gentiohexaose confirms that the preferred glycosyl ligand structure with possible extensions at O1 and O6 extends to longer chains of β-1,6 linked units. These results align well with the previously observed binding of *Cp*CBM92A to pustulan and yeast β-glucan [7].

**Figure 4.**
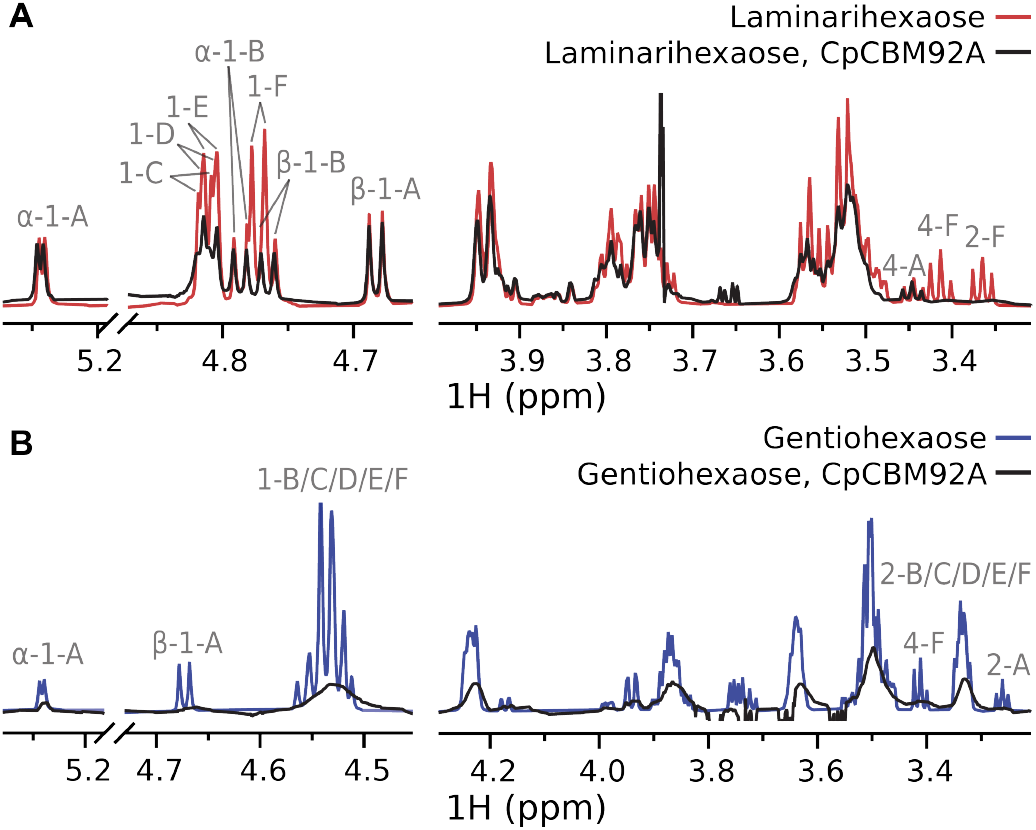
^1^H NMR spectra of laminarihexaose (A) and gentiohexaose (B) in the absence (red/blue) and presence (black) of *Cp*CBM92A, show significant line-broadening of all resonances from gentiohexaose, and resonances on and close to the non-reducing end of laminarihexaose. The spectrum of pure *Cp*CBM92A was subtracted from the ligand spectra in the presence of protein to eliminate interference from protein signals. See ***Figure S14*** and ***Table S1*** for chemical shift assignments.

### Protein-observed experiments reveal different binding modes for each subdomain of *Cp*CBM92A

Because the ligand-observed experiments do not distinguish interactions at the three binding sites of *Cp*CBM92A, we recorded ^1^H, ^15^N-HSQC spectra of the protein in the absence and presence of laminarihexaose and gentiohexaose. These experiments revealed that each binding site plays different roles in binding to β-1,3- and β-1,6-linked oligosaccharides. To visualize the interaction, we supplemented the CSP measurements with predicted protein-ligand complex structures that were generated using Boltz-1x. ***Figure 5*** shows the experimental CSP (see ***Figure S15*** and ***Figure S16***) mapped on the generated structures of *Cp*CBM92A at the three binding sites, α, β, and γ. The predicted structures showed *exo* binding of the non-reducing end of laminarihexaose (***Figure 5***-A), and *endo* binding of gentiohexaose (***Figure 5***-B) at all binding sites, which matches our observations in the ligand-observed experiments (***Figure 4***). For both ligands, we observed strong line-broadening in the NMR spectra at the β binding site (Trp90), reflecting its high affinity to glycosyl units of the preferred structure (***Figure 3***). Interestingly, a different binding mode to laminarihexaose was observed at the α binding site, where a larger surface area of the protein was affected by the presence of the ligand. This subdomain contains two residues, Lys32 and Gln35, that are not present in the β and γ binding sites, but may be involved in polar interactions with longer chains of β-1,3 linked glycosyl units. Similarly, the more distant Thr98 and Asp100 residues on the γ subdomain, which are absent from the other domains, were highly affected by binding to gentiohexaose. This indicates that they may interact with other parts of the β-1,6-linked chain, not only with the glycosyl unit that binds Trp132. With regards to the proposed cross-linking interaction of *Cp*CBM92A with longer polysaccharide ligands, we propose that each binding site contributes differently to this phenomenon. We suggest the following mechanism: the β binding site possibly serves as a primary, high-affinity attachment point, binding for example to theβ-1,6 linked glucosyl substitutions of scleroglucan. Longer chains of β-1,3 linked glycosyl units found between substitution points on scleroglucan or yeast β-glucan bind with lower affinity to the α subdomain, with the non-reducing end stacked with Trp48. The γ binding site may bind longer segments of β-1,6 branches in yeast β-glucan or the non-reducing ends of β-1,3 chains in laminarin or scleroglucan.

**Figure 5.**
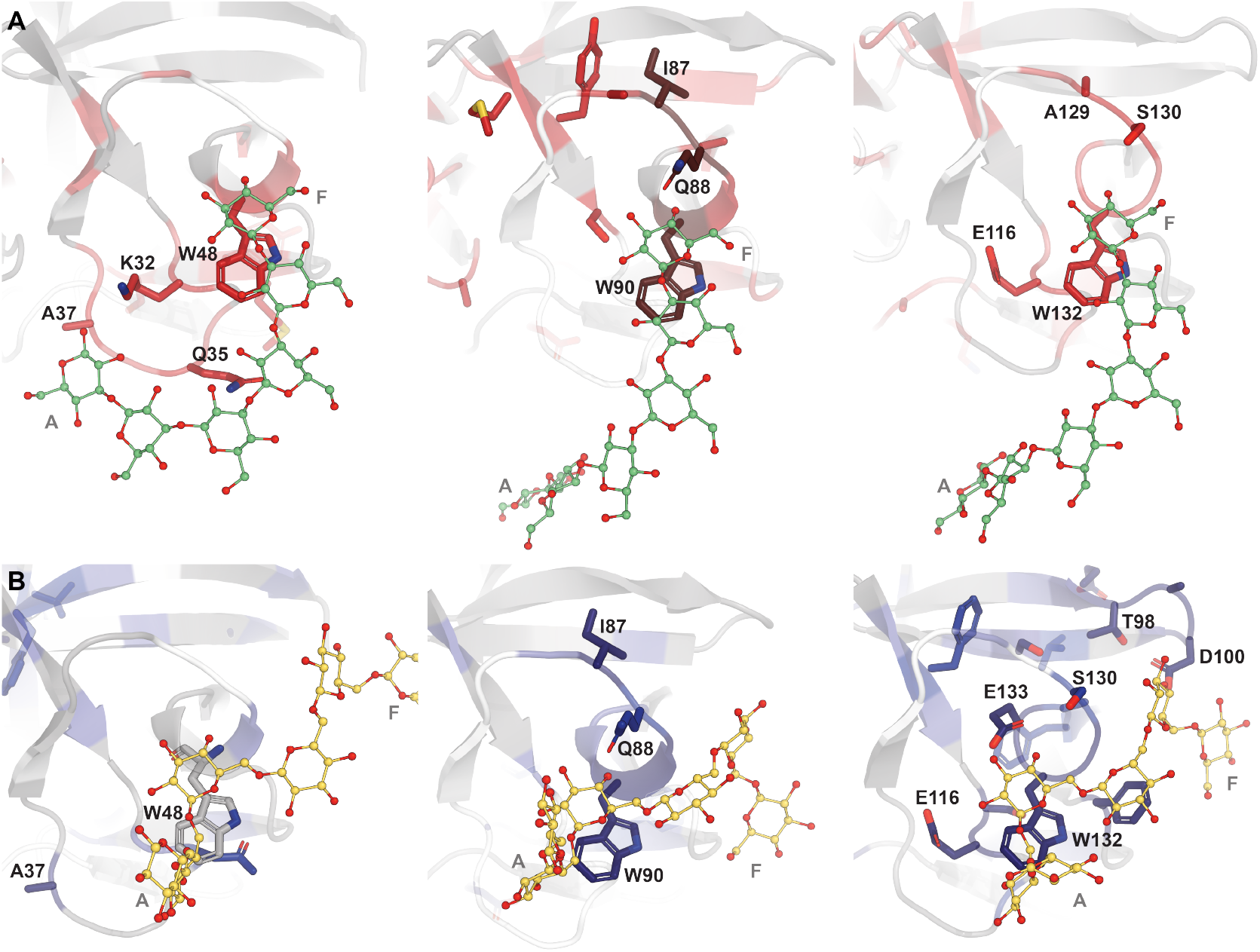
CSP mapping from ^1^H,^15^N-HSQC spectra shown on Boltz-1x predicted structures of *Cp*CBM92A in complex with laminarihexaose (A) and gentiohexaose (B) at the α (left), β (middle) and γ (right) binding sites. Residues with high CSP or high intensity reduction (line-broadening) are shown as stick representations, and the coloring represents the CSPs as shown in ***Figure S15*** and ***Figure S16***. Oligosaccharide ligands are shown as ball and stick structures in green (laminarihexaose) and yellow (gentiohexaose), with labels A and F to mark the reducing and non-reducing ends.

### Binding to laminarin

Laminarin is a useful model substrate to investigate the binding of *Cp*CBM92A to β-1,3-glucans with β-1,6 branches, as it is structurally similar to scleroglucan but has shorter chain length and fewer substitutions (***Figure S17***). Whereas mixing *Cp*CBM92A with scleroglucan led to increases in viscosity that hampered experimentation [15], we did not observe agglutination when incubating laminarin with *Cp*CBM92A, allowing us to perform ligand-observed NMR experiments without signal loss from the formation of a high-molecular-weight complex. We used laminarin from *Laminaria digitata*, and partially assigned the ^1^H chemical shifts based on comparison with laminari- and gentiooligosaccharide spectra (***Figure S18, Table S2***) and previous reports by Kim *et al*. [30]. From the ^1^H NMR spectrum, we estimated an average degree of polymerization (DP_*n*_) of 19 for the β-1,3 linked backbone, with a degree of branching (DB) of about 6%, corresponding to just over one branch per molecule on average. The average branch length was determined to be 1.75, meaning each branch consists of just under two β-1,6 linked glucosyl units on average. The reducing end of the laminarin backbone may consist of a glucosyl (G-series) or mannitol (M-series) residue, and we estimated that about 30% of the residues in our laminarin are G-series. These observations align with previous characterization of laminarin from *Laminaria digitata* [31]. We recorded ^1^H NMR spectra of laminarin in the presence of *Cp*CBM92A, and found that the resonances belonging to protons of the β-1,3-linked backbone (BB) were not affected by the protein (see ***Figure 6***), as both the chemical shifts and linewidths of the corresponding peaks did not differ from what was observed for the free ligand. Similarly, we did not detect a binding effect at the reducing end glucosyl unit (RTg) of the backbone (***Figure 6***-B). The non-reducing end (NRT) displayed line-broadening and CSP for the resonances of protons 1, 2 and 4, which confirms that the NRT of the β-1,3 backbone is interacting with *Cp*CBM92A, as we also observed for the unbranched oligosaccharide laminarihexaose (***Figure 4***-A). According to our results on gentiohexaose (***Figure 4***-B), *endo* binding of *Cp*CBM92A along a β-1,6 branch should be possible. For laminarin, we observed line broadening and CSP only at the terminal glucosyl units of the branches (SCT) (***Figure 6***-C-F). This indicates that *Cp*CBM92A prefers to bind *exo* to short β-1,6 branches consisting of one or two glucosyl units, as the presence of the polysaccharide backbone prevents bidning to non-terminal glucosyl units, suggesting that laminarin and scleroglucan will be bound only in the *exo* mode at their substitution points. However, polysaccharides containing longer β-1,6 branches, like yeast β-glucan, or polymeric linear β-1,6 glucans (pustulan) could still be acceptable substrates for *endo* binding.

**Figure 6.**
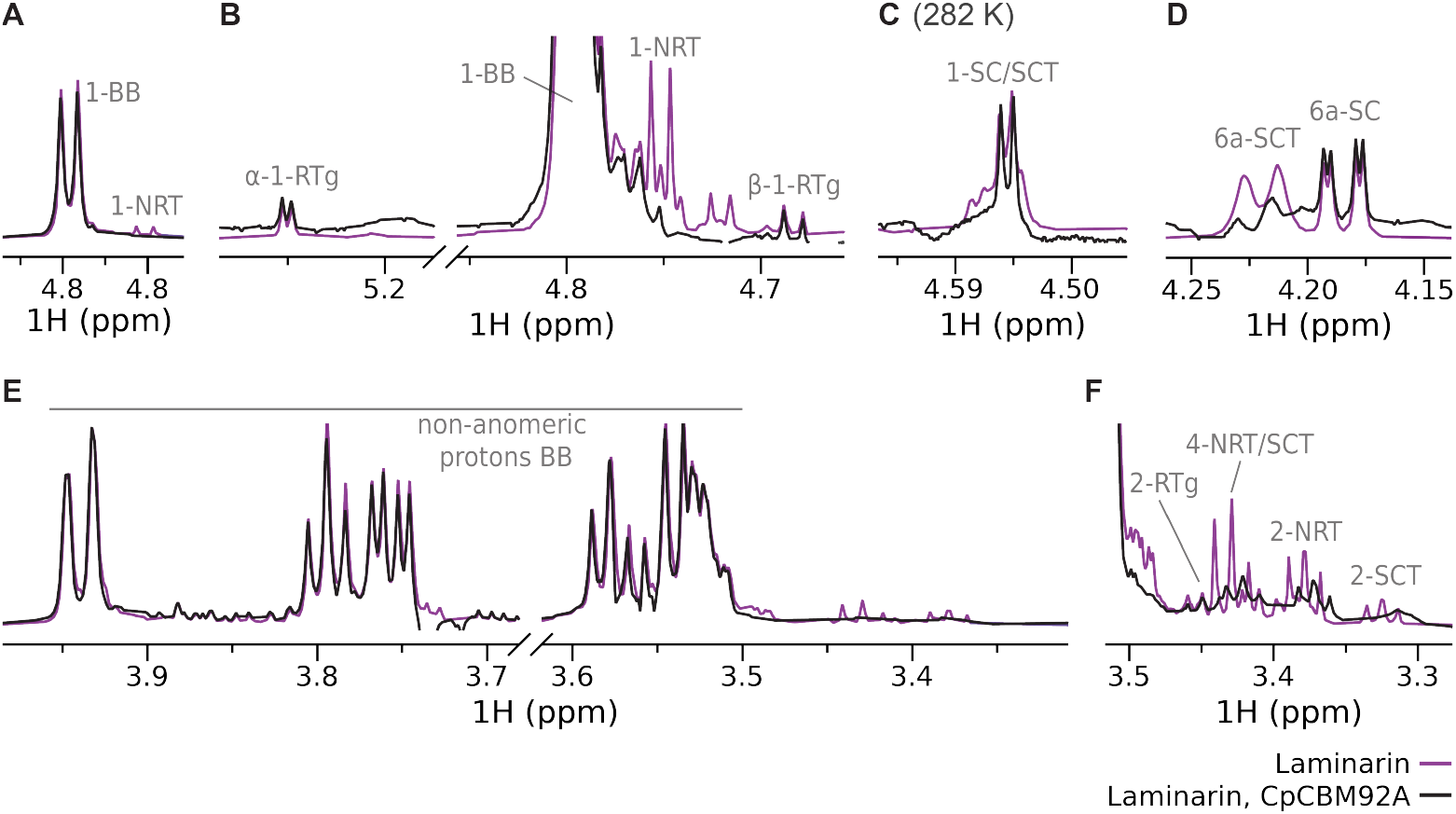
^1^H NMR spectra of laminarin in the absence (purple) and presence (black) of *Cp*CBM92A, at 318 K (A, B, D-F) and 282 K (C). The spectrum of pure *Cp*CBM92A was subtracted from the ligand spectrum in the presence of protein to eliminate interference from protein signals. Relevant assignments are shown on the spectra, BB = β-1,3 backbone, RTg = reducing end of G-series β-1,3 backbone, NRT = non-reducing end of β-1,3 backbone, SC = β-1,6 side chain, SCT = non-reducing end of β-1,6 side chain. The numbers correspond to the proton numbering on the glucose rings (***Figure S17***). Full spectra are shown in ***Figure S18***.

## Methods and Materials

### Carbohydrate substrates

Glucose, cellobiose (219458), and laminarin (L9634) were purchased from Merck (Germany). Gentiobiose (OG05175) and laminaribiose (OL02452) were purchased from Biosynth (UK). Sophorose (O-SOPH) and laminarihexaose (O-LAM6) were purchased from Megazyme (Ireland). Pustulan (GLU900) was purchased from Elicityl (France).

### Production of gentiohexaose

Gentiohexaose was produced in-house by partial acid hydrolysis of pustulan. Pustulan (100 mg) was dissolved in 50 mL aqueous NaCl (50 mM) overnight. HCl was added to a final concentration of 0.01 M, and the sample was diluted to a final volume of 100 mL with 50 mM NaCl solution. The solution was incubated at 95°C with gentle shaking for 49.5 h, then cooled to rt and neutralized with NaOH. The sample was freeze-dried, and the residue was dissolved in 5 mL MQ water and filtered through a 0.45 µm filter. Gentiooligosaccharides were separated by semi-preparative SEC by injecting 2 mL of the solution onto three serially connected HiLoad 26/600 Superdex 30 columns (Cytiva, USA), with MQ water as the eluent and a flow rate of 0.8 mL min/min (see ***Figure S19***). Gentiohexaose was isolated in 6.3% yield (2.5 mg). The degree of polymerization of the oligosaccharides was confirmed by NMR.

### Protein production and purification

The plasmid for expression of *Cp*CBM92A (with C-terminal His_6_-tag) was constructed as described previously [7]. Chemically competent *E. coli* BL21 (DE3) cells (New England Biolabs, USA) were transformed with the plasmid by heat-shock, and grown in 5 mL LB pre-cultures supplemented with 100 µg/mL ampicillin at 37°C and 225 rpm overnight. For production of isotopically labelled ^15^N,^13^C-*Cp*CBM92A and ^15^N-*Cp*CBM92A, 5 mL pre-culture was used to inoculate main cultures of 500 mL M9 media (6 g/L Na_2_HPO_4_, 3 g/L KH_2_PO, 0.5 g/L NaCl) supplemented with 2 mM MgSO_4_, 0.1 mM CaCl_2_, 10 mL BioExpress Cell Growth Media (Cambridge Isotope Laboratories, USA), 5 mL Gibco™ MEM Vitamin Solution (100x, Thermo Fisher Scientific, USA), 500 µL trace metal solution (1000x, containing 10 g/L CaCl_2_, 8 g/L MnSO_4_, 5 g/L FeSO_4_, 1 g/L CuSO_4_, and 1 g/L ZnSO_4_), 100 µg/mL ampicillin, 1 g/L (^15^NH_4_)_2_SO_4_ and either 4 g/L ^13^C-glucose or 1% natural abundance glycerol. For production of non-labelled *Cp*CBM92A, 5 mL pre-culture was used to inoculate main cultures of 500 mL 2xLB supplemented with 100 µg/mL ampicillin. The main cultures were incubated in a LEX bioreactor (Epiphyte3, Canada) at 22°C. At OD_600_ = 1.8-2.0, the production of *Cp*CBM92A was induced by adding 0.1 mM isopropyl-β-D-1-thiogalactopyranoside (IPTG) and the culture was incubated overnight at 22°C.

Cells were harvested by centrifugation at 4°C and 5000g for 10 min. The cell pellet was gently resuspended in 25 mL 50 mM Tris-HCl (pH 8.0) containing 300 mM NaCl and one tablet of EDTA-free protease inhibitor (Roche), and lysed by sonication for 10 min (cycle of 5 s on, 10 s off), followed by centrifugation at 4°C and 25000g for 30 min. The lysate was filtered through a 0.2 µm filter and the protein was purified using a 1 mL HisTrap™ HP Ni affinity column (Cytiva) on an ÄKTApure FPLC system (Cytiva), at a flow rate of 1 mL/min. The protein was eluted with an imidazole gradient (0-400 mM) in 50 mM Tris-HCl (pH 8.0) containing 300 mM NaCl (see ***Figure S20***). The fractions were analyzed with SDS-PAGE using 12% SurePAGE Bis-Tris gels (GenScript), using PAGE-MASTER Protein Standard Plus (GenScript) for the identification of the target protein (see ***Figure S21***). Fractions containing *Cp*CBM92A were pooled, concentrated using a Vivaspin® concentrator (Sartorius) with a 10 kDa cut-off, and the buffer was exchanged to the appropriate buffer for each NMR experiment using a PD MidiTrap G-25 column (Cytiva). Protein concentration was determined by measuring A280 on a NanoDrop ND-1000 spectrophotometer (Thermo Fisher Scientific) with the theoretical extinction coefficient (ε = 37470 M^−1^cm^−1^), calculated using the ProtParam tool (web.expasy.org/protparam/) [32]. The yields of isolated protein were 3.2 mg (^15^N,^13^C-*Cp*CBM92A), 30 mg (^15^N-*Cp*CBM92A) and 90 mg (*Cp*CBM92A) per L of culture.

### NMR experiments

NMR spectra were recorded on a Bruker Ascend 800 MHz NMR spectrometer (Bruker BioSpin AG, Switzerland) equipped with an Avance NEO console and a 5 mm cryogenic CP-TCI probe, at the NV-NMR Center at NTNU, which is part of the Norwegian NMR Platform (NNP). NMR spectra were recorded and processed in Bruker TopSpin version 4.4.0. 1D spectra were further analysed and visualised in MestReNova version 14.1.12. Annotation of assignments on the spectra was done in Adobe Illustrator 2025.

#### Backbone chemical shift assignment of *Cp*CBM92A

For the backbone assignment of *Cp*CBM92A, the following NMR spectra were acquired at 298 K: ^1^H,^15^N-HSQC, HNCA (BEST-TROSY), HNcoCACB ((BEST-TROSY), HNCACO ((BEST-TROSY), HNCACB ((BEST-TROSY), HBHAcoNH, ^15^N-NOESY, G-(i+1)-HSQC (MUSIC, ^1^H,^15^N-HSQC, Gly+1 residues [19]) and W-(NHε)-HSQC (MUSIC, ^1^H,^15^N-HSQC, Trp sidechains [20]). NMR spectra were analysed using CcpNmr AnalysisAssign [33]). The chemical shift assignment of backbone resonances of *Cp*CBM92A (>84%) is reported here. The NMR sample used for these experiments contained purified ^15^N,^13^C-*Cp*CBM92A (70 µM) in 20 mM MES (pH 6.0) with 10 mM NaCl, supplemented with 10% D_2_O.

#### Protein-observed titrations with disaccharides

CSP of ^15^N-*Cp*CBM92 amide groups were measured by recording ^15^N-HSQC spectra at 298 K for varying concentrations of glucose and disaccharide ligands. A reference spectrum was recorded for a sample of ^15^N-*Cp*CBM92 (100 µM) in assay buffer (25 mM sodium phosphate (pH 6.5) containing 10 mM NaCl, supplemented with 10% D_2_O). Glucose (16.15 mM), sophorose (8.26 mM), laminaribiose (9.91 mM), cellobiose (9.91 mM) and gentiobiose (4.20 mM) were dissolved in separate samples containing ^15^N-*Cp*CBM92 (100 µM) in assay buffer, and the samples were then combined with the reference sample in appropriate proportions to obtain samples of 5 intermediate concentrations. ^1^H,^15^N-HSQC spectra were recorded for each concentration. The CSP Δδ was calculated as 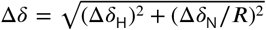, where Δδ_H_ and Δδ_N_ are the CSP of the amide proton and nitrogen in ppm, and the chemical shift scaling factor *R* was set to 6.5 [34]. The dissociation constant *K*_*D*_ was determined by fitting the CSP to a two-site fast exchange model,

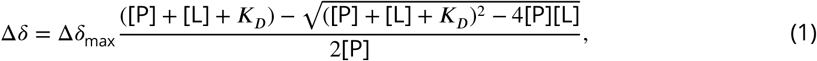

where Δδ_max_ is the CSP at saturation, [P] is the protein concentration, and [L] is the ligand concentration. The CSP analysis was done using CcpNmr AnalysisAssign [33]).

#### Protein-observed chemical shift mapping with oligosaccharides

CSP of ^15^N-*Cp*CBM92 amide groups upon addition of oligosaccharide ligands was measured by recording ^1^H,^15^N-HSQC spectra at 298 K for 80 µM ^15^N-*Cp*CBM92 in assay buffer. New spectra were recorded for samples containing 800 µM laminarihexaose or gentiohexaose. The CSP were calculated as described above.

#### Ligand-observed experiments

For each ligand-observed NMR experiment, two spectra were recorded for each ligand; one for a reference sample containing only 340 µM glucose, 500 µM laminaribiose/gentiobiose, 800 µM laminarihexaose/gentiohexaose, or 2.7 mg/mL laminarin in assay buffer supplemented with TSP (150 µM), and one for a sample with 17 µM (glucose sample), 25 µM (laminaribiose/gentiobiose sample), 80 µM (laminarihexaose/gentiohexaose sample), or 100 µM (laminarin sample) *Cp*CBM92A or ^15^N-*Cp*CBM92A added. Spectra were recorded at 298 K for the glucose and disaccharide samples, 282 K for the hexaose and laminarin samples, and 318 K for the laminarin sample. Spectra were also acquired for reference samples of *Cp*CBM92A in the specified concentrations to subtract from the ligand spectra in the presence of protein to eliminate protein signals. ^1^H NMR experiments were recorded using a standard pulse sequence with NOESY presaturation water suppression, with 65k points and a spectral width of 12 ppm. WaterLOGSY experiments were recorded using a 1D Water-LOGSY pulse sequence with excitation sculpting water suppression, and a 50 ms spin-lock to partly suppress protein signals (ephogsygpno.2) [35]. The WaterLOGSY mixing time was 1.5 s. Spectra were recorded with 65k points and 256 scans. The WaterLOGSY factor was calculated as 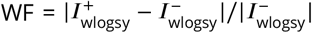, where 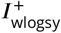 and 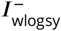 are the intensities of the proton peaks in the WaterLOGSY spectrum in the presence and absence of protein [22]. Because of some peak overlap of multiplets in the ^1^H spectra, 1-4 distinguishable lines were picked for each proton (see ***Figure S8, Figure S9, Figure S10***), and their WF were averaged to give an overall WF, and the error was calculated as their standard deviation. Due to CSP, the WaterLOGSY spectra were not subtracted before peak picking to calculate 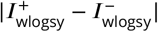. Instead, distinguishable peaks were picked in both spectra, and 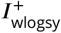 and 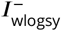 were determined separately.

### Structure predictions

The structures of *Cp*CBM92A-ligand complexes were predicted using Boltz-1x [16, 24]. The protein sequence was retrieved from [7] (PBD ID: 7ZOI), and the ligands were provided as SMILES strings. Glucose and disaccharide ligands were re-docked to the binding sites identified by Boltz-1x as an additional step using Gnina [28]. We performed 20 dockings per ligand in Gnina. Predicted *K*_*d*_ values were calculated by averaging the CNN affinities for all structures with a CNN score above 0.85.

## Data availability

Data and scripts used for analysis and for generating the figures are available from https://github.com/gcourtade/papers/blob/master/2026/CBM92/

## Acknowledgments

G.C. gratefully acknowledges funding by the Novo Nordisk Foundation (grant number NNF22OC0073963). L.S.M. acknowledges support from the Swedish Research Council Formas (grant 2019-00389).

