## Supplementary information for "Molecular basis of protein-glycan cross-linking by *Cp*CBM92A revealed by NMR spectroscopy"

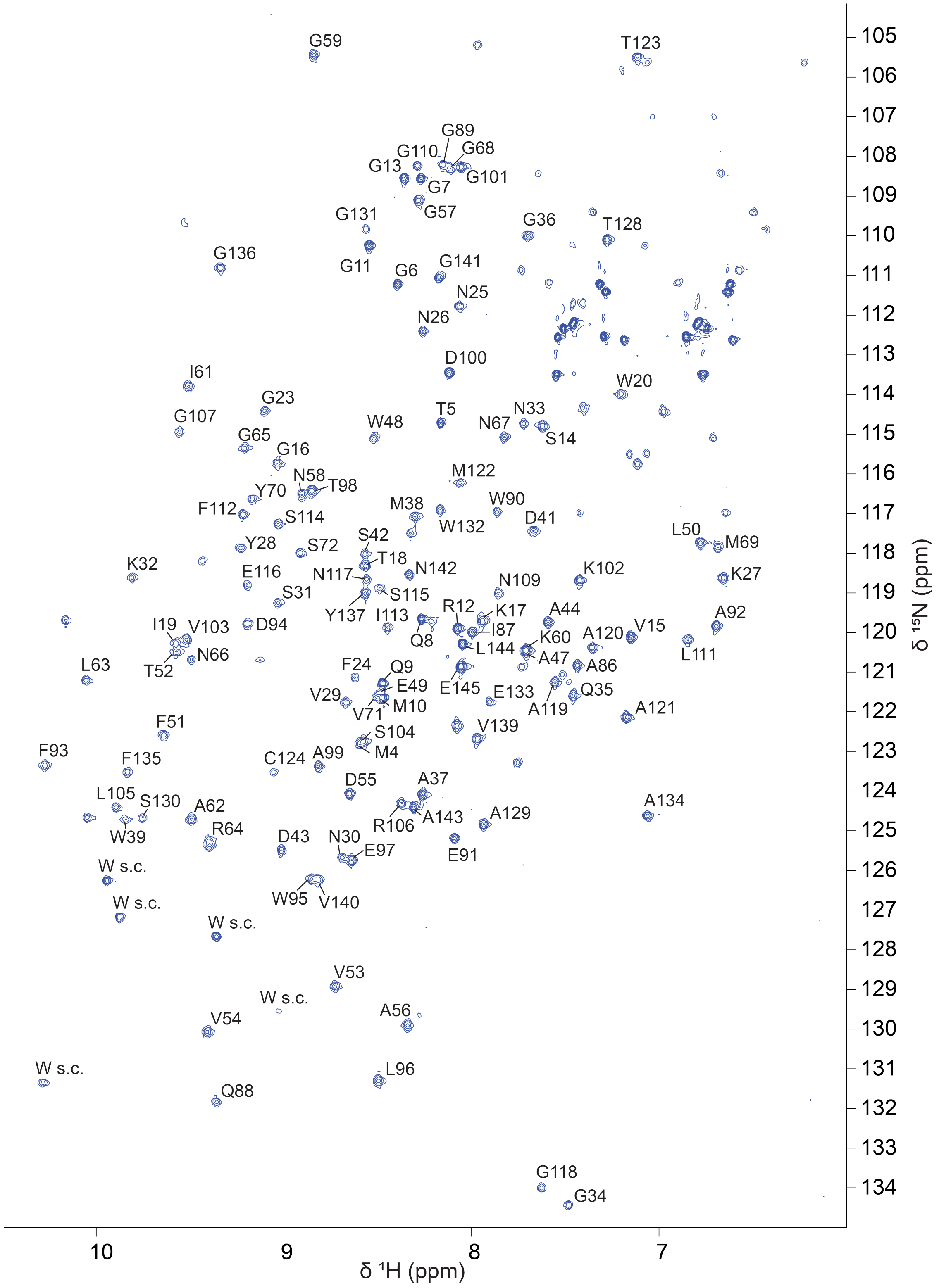


**Figure S1.** $^{1}H,^{15}N$-HSQC spectrum of $^{1}H,^{15}N,^{13}C$-labeled *Cp*CBM92A in 20 mM MES (pH6.0) supplemented with 10 mM NaCl and 10% $D_2O$, with assignment of resonances to residues in the protein sequence. >84% of the residues were assigned. Unassigned Trp side chains are labeled as W s.c.. Sequence: MAS**MTGGQQMGRGSVGKTIW**LQ**GFNNKYVNSKNGQGAMW**C**DSDA**PQ**AWELFTVVDAGNGKIALRGNNGMYVS**SENGEQAITCNRP**AIQGWEAFDWLETADGKVSLRG**S**NGLFISSENGAAAMTC**TRP**TASGWEAFGY**S**VVGNALE**HHHHHH (assigned residues are highlighted in bold).

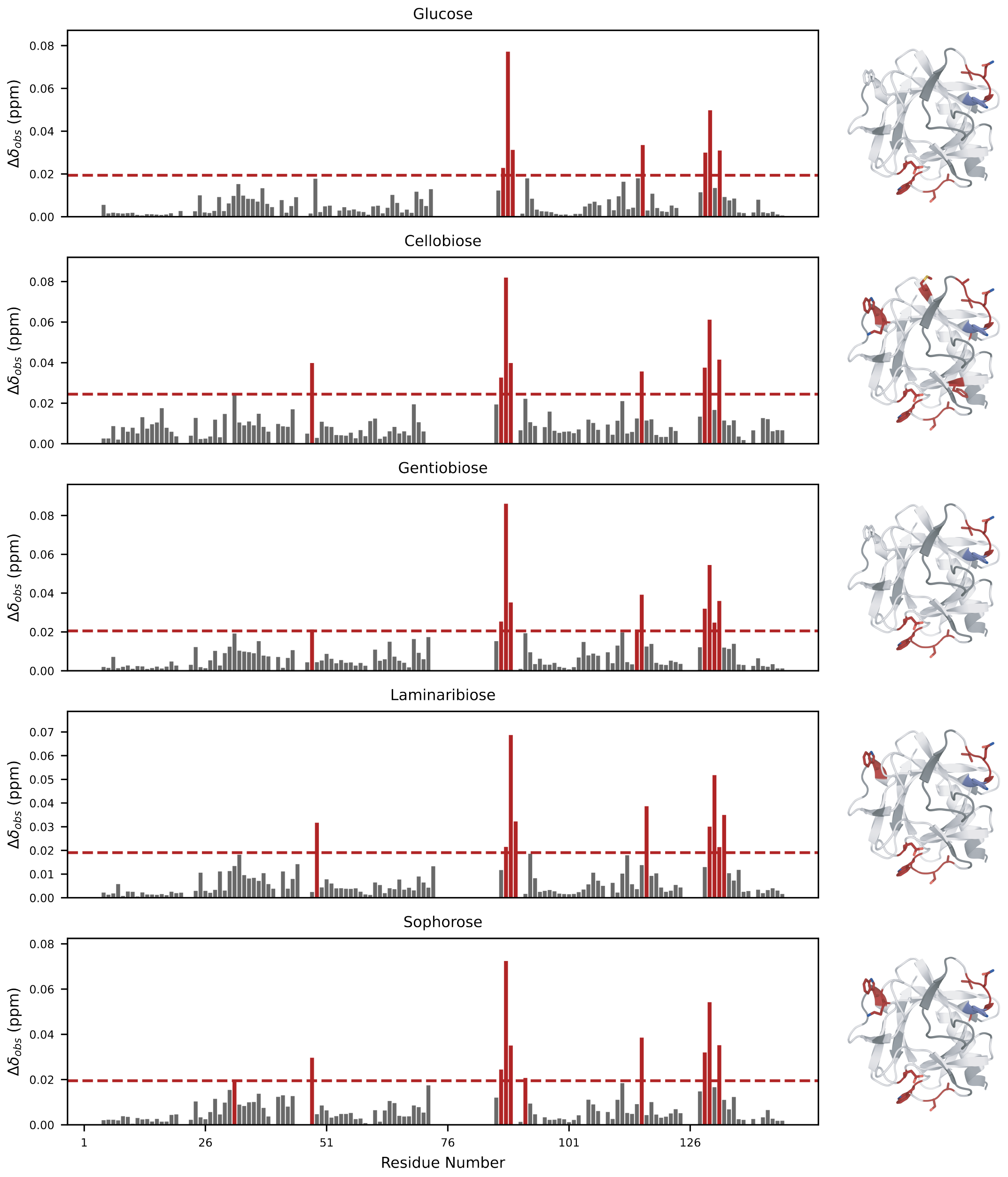


**Figure S2.** Chemical shift perturbations (CSP) $\Delta\delta$ in ppm measured for $^{1}H$,$^{15}N$-labeled *Cp*CBM92A during titrations with glucose, cellobiose, gentiobiose, laminaribiose and sophorose. High CSP (one standard deviation above the mean CSP value) are indicated in red on the charts and structure. Trp 90 is colored blue to illustrate high intensity reduction, which was observed instead of CSP for this residue. Unassigned residues are colored in grey.

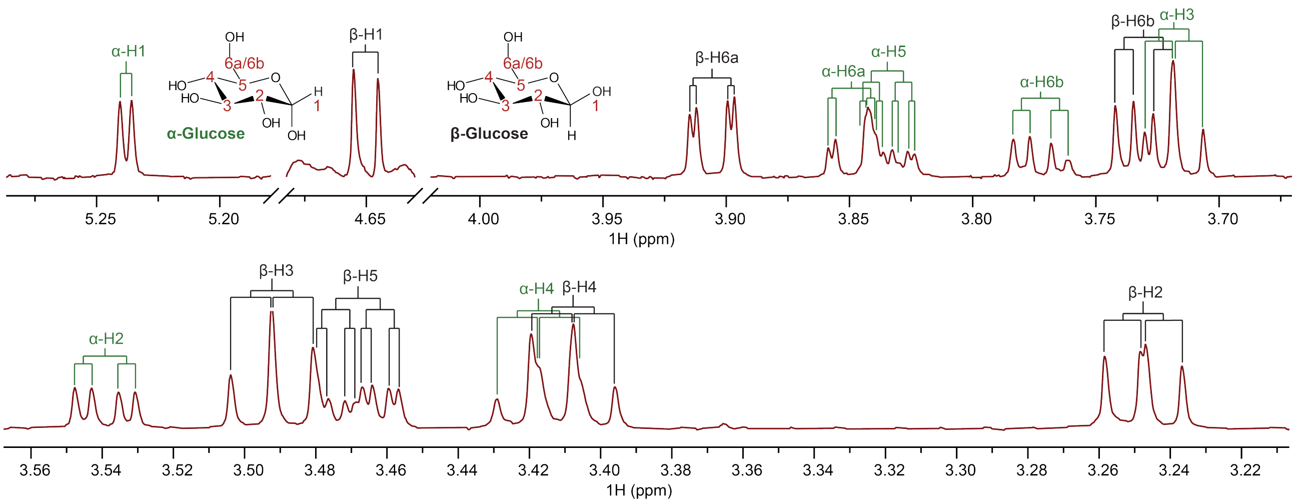


**Figure S3.** $^{1}$H NMR spectrum (800 MHz) of glucose acquired at 298 K. The spectrum is calibrated based on the TSP reference signal at 0 ppm.

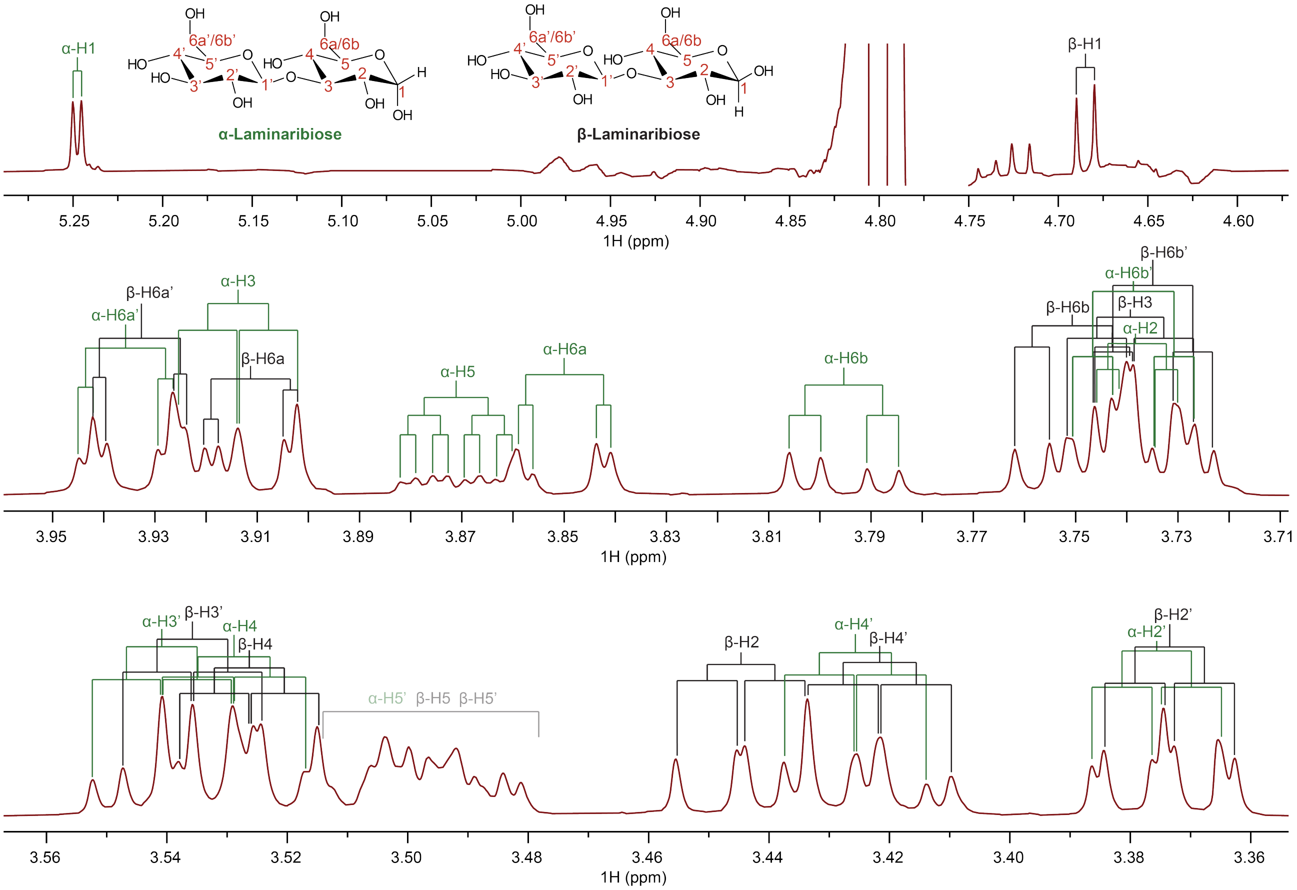


**Figure S4.** $^{1}$H NMR spectrum (800 MHz) of laminaribiose acquired at 298 K. The spectrum is calibrated based on the TSP reference signal at 0 ppm.

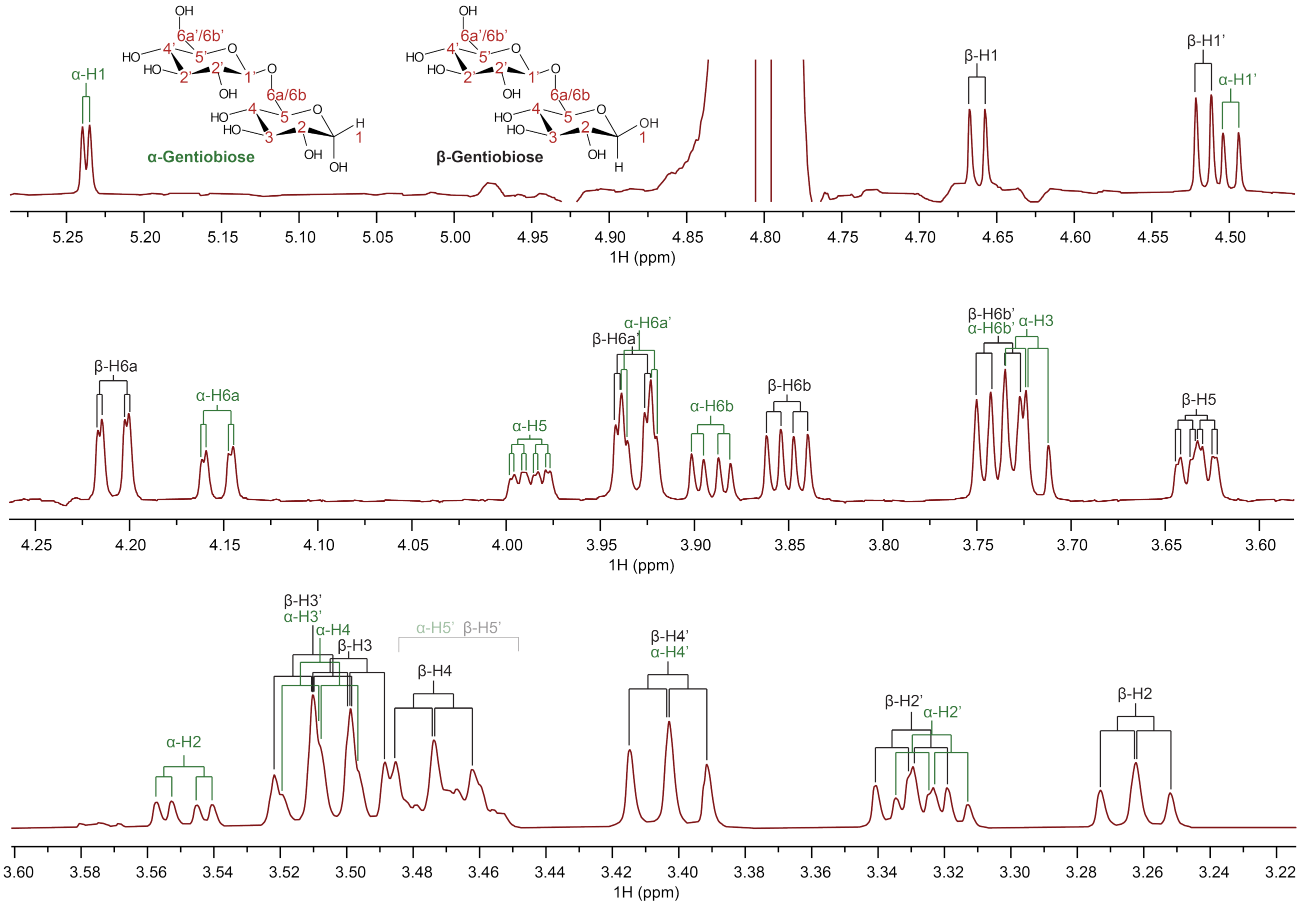

**Figure S5.** $^{1}H$ NMR spectrum (800 MHz) of gentiobiose acquired at 298 K. The spectrum is calibrated based on the TSP reference signal at 0 ppm.

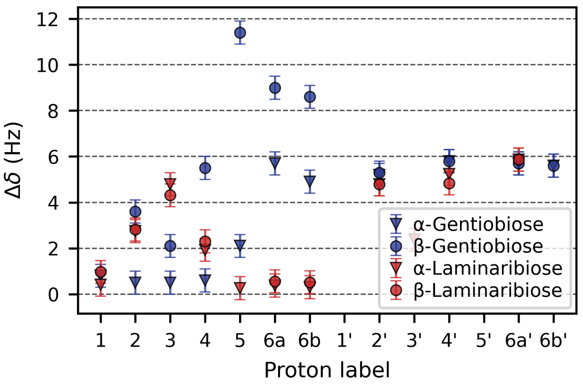


**Figure S6.** Chemical shift perturbations (CSP) Δ$\delta$ in Hz measured for 500 μM laminaribiose and gentiobiose upon addition of 25 μM *Cp*CBM92A. Error bars represent the estimated maximum error (0.5 Hz) for manual measurement of CSP based on the peak widths.

**Table S1.** $^{1}$H chemical shift (δ) assignments and coupling constants ($J$) for *Cp*CBM92A mono- and disaccharide ligands, from NMR spectra acquired at 298 K. The proton labels are as shown in ***Figure S3***, ***Figure S4*** and ***Figure S5***. Chemical shifts labelled with an asterisk (*) indicate overlapping peaks where the individual chemical shifts were indistinguishable. The relative amounts of α and β anomers (calculated from integration of anomeric proton signals) are indicated as percentages for each ligand.

| **α-Glucose** (47%) | | | **β-Glucose** (53%) | | |
|---|---|---|---|---|---|
| Proton | δ (ppm) | $J$(H$_i$-H$_{i+1}$) (Hz) | Proton | δ (ppm) | $J$(H$_i$-H$_{i+1}$) (Hz) |
| 1 | 5.238 | 3.7 | 1 | 4.650 | 7.8 |
| 2 | 3.539 | 9.8 | 2 | 3.247 | 9.2 |
| 3 | 3.718 | 9.6 | 3 | 3.492 | 9.3 |
| 4 | 3.417 | 9.5 | 4 | 3.408 | 9.4 |
| 5 | 3.834 | 2.3 (6a), 5.3 (6b) | 5 | 3.468 | 2.2 (6a), 5.9 (6b) |
| 6a | 3.849 | 12.2 (6b) | 6a | 3.906 | 12.2 (6b) |
| 6b | 3.772 | | 6b | 3.730 | |
| **α-Laminaribiose** (52%) | | | **β-Laminaribiose** (48%) | | |
| Proton | δ (ppm) | $J$(H$_i$-H$_{i+1}$) (Hz) | Proton | δ (ppm) | $J$(H$_i$-H$_{i+1}$) (Hz) |
| 1 | 5.248 | 3.8 | 1 | 4.685 | 8.1 |
| 2 | 3.736 | 9.3 | 2 | 3.445 | 9.2 |
| 3 | 3.914 | 9.1 | 3 | 3.739 | 9.1 |
| 4 | 3.529 | 9.9 | 4 | 3.526 | 9.9 |
| 5 | 3.871 | 2.3 (6a), 5.0 (6b) | 5 | (3.515-3.478)* | 2.2 (6a), 5.6 (6b) |
| 6a | 3.850 | 12.3 (6b) | 6a | 3.912 | 12.4 |
| 6b | 3.795 | | 6b | 3.751 | |
| 1′ | | 9.0 | 1′ | | 8.0 |
| 2′ | 3.375 | 9.5 | 2′ | 3.373 | 9.4 |
| 3′ | 3.541 | 9.2 | 3′ | 3.536 | 9.1 |
| 4′ | 3.426 | 9.8 | 4′ | 3.422 | 9.9 |
| 5′ | (3.515-3.478)* | 2.1 (6a′), 6.3 (6b′) | 5′ | (3.515-3.478)* | 2.0 (6a′), 6.2 (6b′) |
| 6a′ | 3.936 | 12.3 (6b′) | 6a′ | 3.933 | 12.3 (6b′) |
| 6b′ | 3.737 | | 6b′ | 3.735 | |
| **α-Gentiobiose** (39%) | | | **β-Gentiobiose** (61%) | | |
| Proton | δ (ppm) | $J$(H$_i$-H$_{i+1}$) (Hz) | Proton | δ (ppm) | $J$(H$_i$-H$_{i+1}$) (Hz) |
| 1 | 5.237 | 3.7 | 1 | 4.662 | 7.9 |
| 2 | 5.549 | 9.8 | 2 | 3.262 | 8.4 |
| 3 | 3.720 | 9.5 | 3 | 3.499 | 9.2 |
| 4 | 3.508 | 10.1 | 4 | 3.474 | 8.5 |
| 5 | 3.987 | 2.1 (6a), 5.1 (6b) | 5 | 3.633 | 2.0 (6a), 5.9 (6b) |
| 6a | 4.153 | 11.5 (6b) | 6a | 4.209 | 11.5 (6b) |
| 6b | 3.891 | | 6b | 3.851 | |
| 1′ | 4.499 | 8.1 | 1′ | 4.517 | 8.0 |
| 2′ | 2.323 | 9.5 | 2′ | 3.330 | 9.5 |
| 3′ | 3.510 | 9.5 | 3′ | 3.510 | 9.5 |
| 4′ | 3.403 | 9.5 | 4′ | 3.403 | 9.5 |
| 5′ | 3.465 | 2.4 (6a′), 6.2 (6b′) | 5′ | 3.465 | 2.3 (6a′), 6.2 (6b′) |
| 6a′ | 3.929 | 12.2 (6b′) | 6a′ | 3.933 | 12.2 (6b′) |
| 6b′ | 3.739 | | 6b′ | 3.739 | |

**Table S2.** Partial [1]H chemical shift (δ) assignments and coupling constants (*J*) for *Cp*CBM92A oligosaccharide ligands. The proton labels are as shown in ***Figure S14*** and ***Figure S17***. The relative amounts of α and β anomers of the reducing ends (A) (calculated from integration of anomeric proton signals), and the temperatures during acquisition are indicated in parentheses.

| **Laminarihexaose** (43% α, 57% β, 282 K) | | | | | |
|---|---|---|---|---|---|
| Proton | δ (ppm) | $J(H_i\text{-}H_{i+1})$ (Hz) | Proton | δ (ppm) | $J(H_i\text{-}H_{i+1})$ (Hz) |
| 1-A (α) | 5.242 | 3.7 | 1-E | 4.809 | 8.0 |
| 1-A (β) | 4.683 | 8.0 | 1-F | 4.772 | 8.0 |
| 1-B (α) | 4.786 | 8.1 | 2-A | 3.444 | 8.9 |
| 1-B (β) | 4.764 | 8.1 | 2-F | 3.365 | 9.1 |
| 1-C | 4.814 | 8.0 | 4-F | 3.413 | 9.5 |
| 1-D | 4.811 | 8.0 | | | |
| **Gentiohexaose** (42% α, 58% β, 282 K) | | | | | |
| Proton | δ (ppm) | $J(H_i\text{-}H_{i+1})$ (Hz) | Proton | δ (ppm) | $J(H_i\text{-}H_{i+1})$ (Hz) |
| 1-A (α) | 5.241 | 3.7 | 6a-F | 3.940 | 12.2 (6b-F), 1.5 (5-F) |
| 1-A (β) | 4.673 | 8.2 | 2-A (α) | 3.546 | 9.6 |
| 1-B/C/D/E/F | 4.573-4.505 | | 4-F | 3.411 | 9.4 |
| 6a-A/B/C/D/E | 4.263-4.204 | | 2-B/C/D/E/F | 3.357-3.311 | |
| 6a-A (α) | 4.171 | 11.1 (6b-A), 1.5 (5-A) | 2-A (β) | 3.261 | 8.5 |
| **Laminarin** (318 K) | | | | | |
| Proton | δ (ppm) | $J(H_i\text{-}H_{i+1})$ (Hz) | Proton | δ (ppm) | $J(H_i\text{-}H_{i+1})$ (Hz) |
| 1-RTg (α) | 5.250 | 3.8 | 6a-SC | 4.185 | 11.0 (6b-SC), 2.5 (5-SC) |
| 1-BB | 4.796 | 7.9 | 2-RTg | 3.450 | 8.0 |
| 1-NRT | 4.751 | 7.9 | 4-NRT/SCT | 3.429 | 9.4 |
| 1-RTg (β) | 4.683 | 7.9 | 2-NRT | 3.378 | 8.5 |
| 1-SC/SCT | 4.590-4.518 | | 2-SCT | 3.324 | 9.5 |
| 6a-SCT | 4.220 | 11.7 (6b-SCT), - | | | |

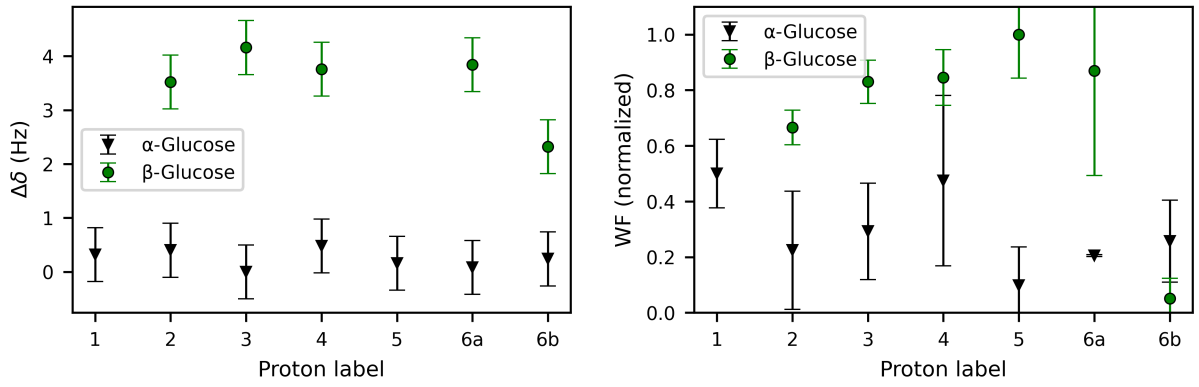


**Figure S7.** Chemical shift perturbations (CSP) $\Delta\delta$ in Hz (left) and WaterLOGSY factors WF (right) measured for 340 µM glucose upon addition of 17 µM *Cp*CBM92A. Error bars represent the estimated maximum error (0.5 Hz) for manual measurement of CSP based on the peak widths (left) and the standard deviation of WFs calculated for each line picked for a specific proton (see ***Figure S8***) (right).

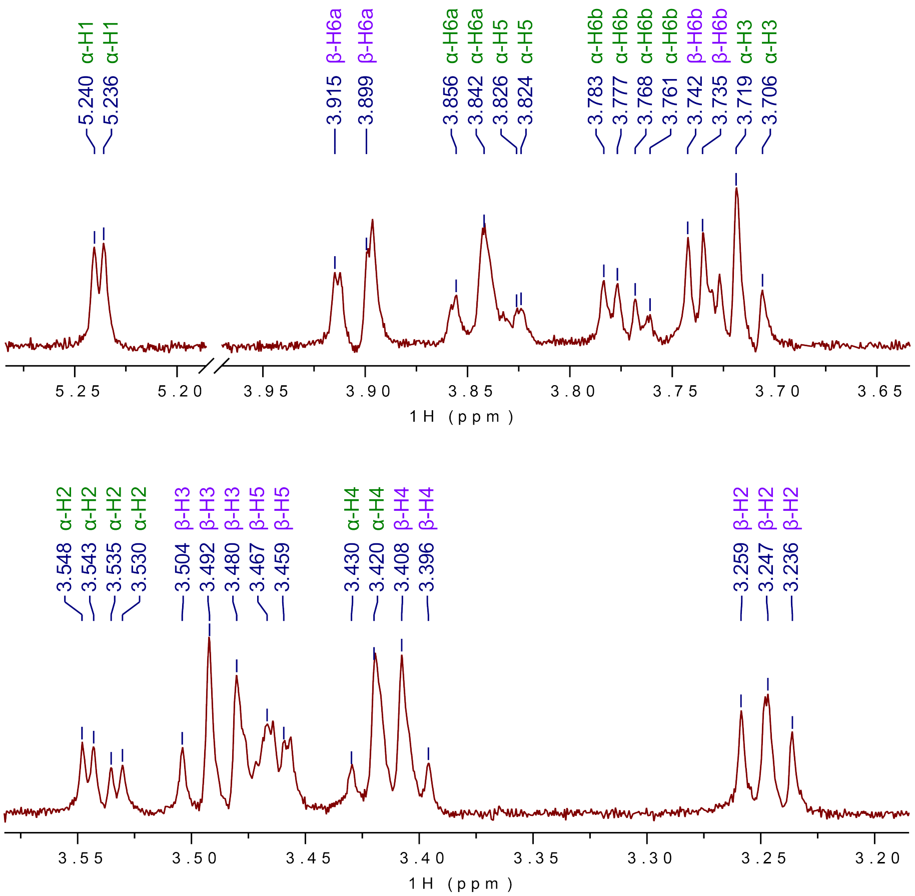


**Figure S8.** $^{1}H$ NMR spectrum of glucose, showing the individual lines picked to determine WaterLOGSY factors (WF).

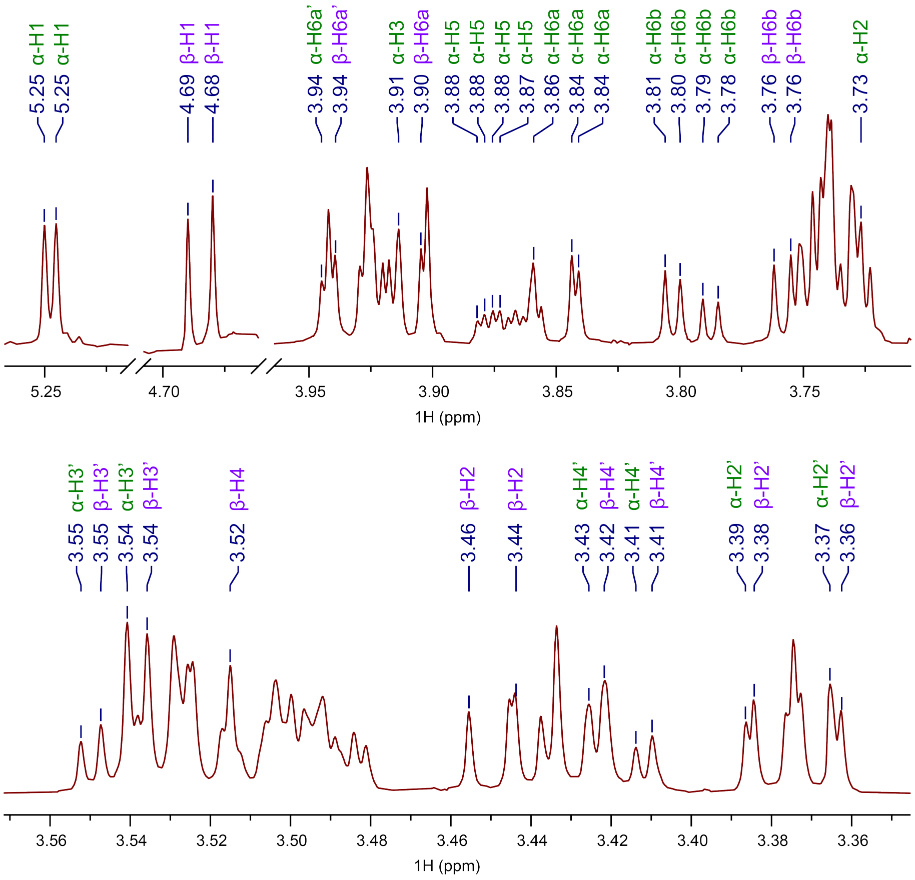


**Figure S9.** $^{1}H$ NMR spectrum of laminaribiose, showing the individual lines picked to determine WaterLOGSY factors (WF).

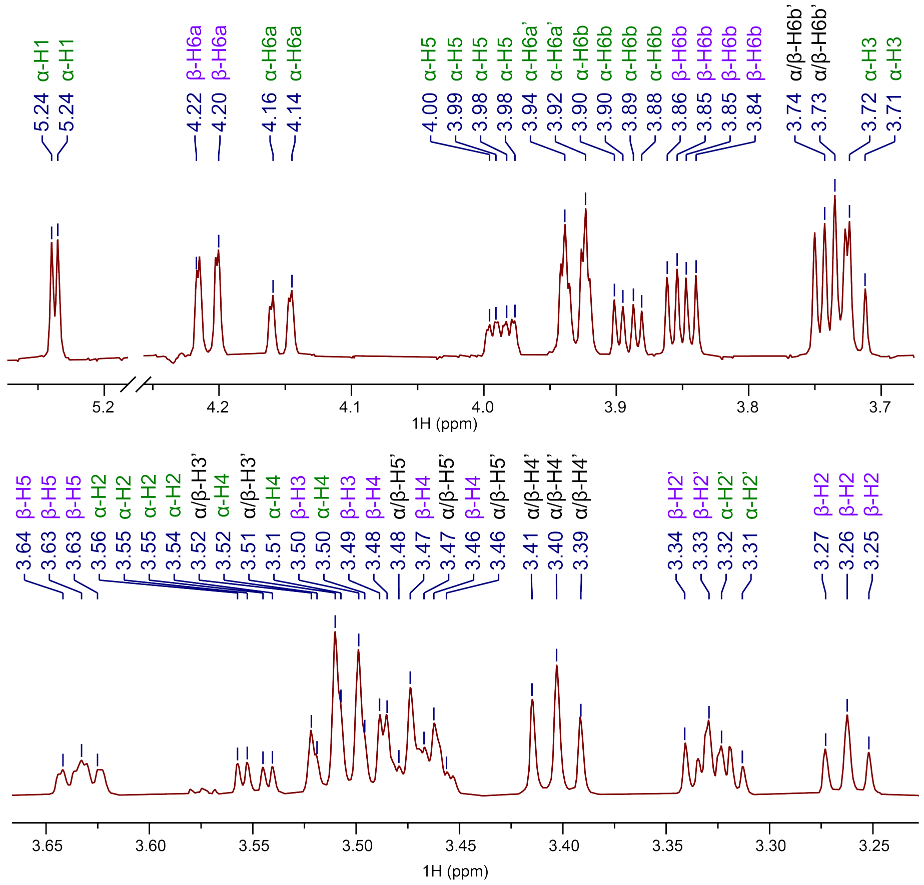


**Figure S10.** $^{1}$H NMR spectrum of gentiobiose, showing the individual lines picked to determine WaterLOGSY factors (WF).

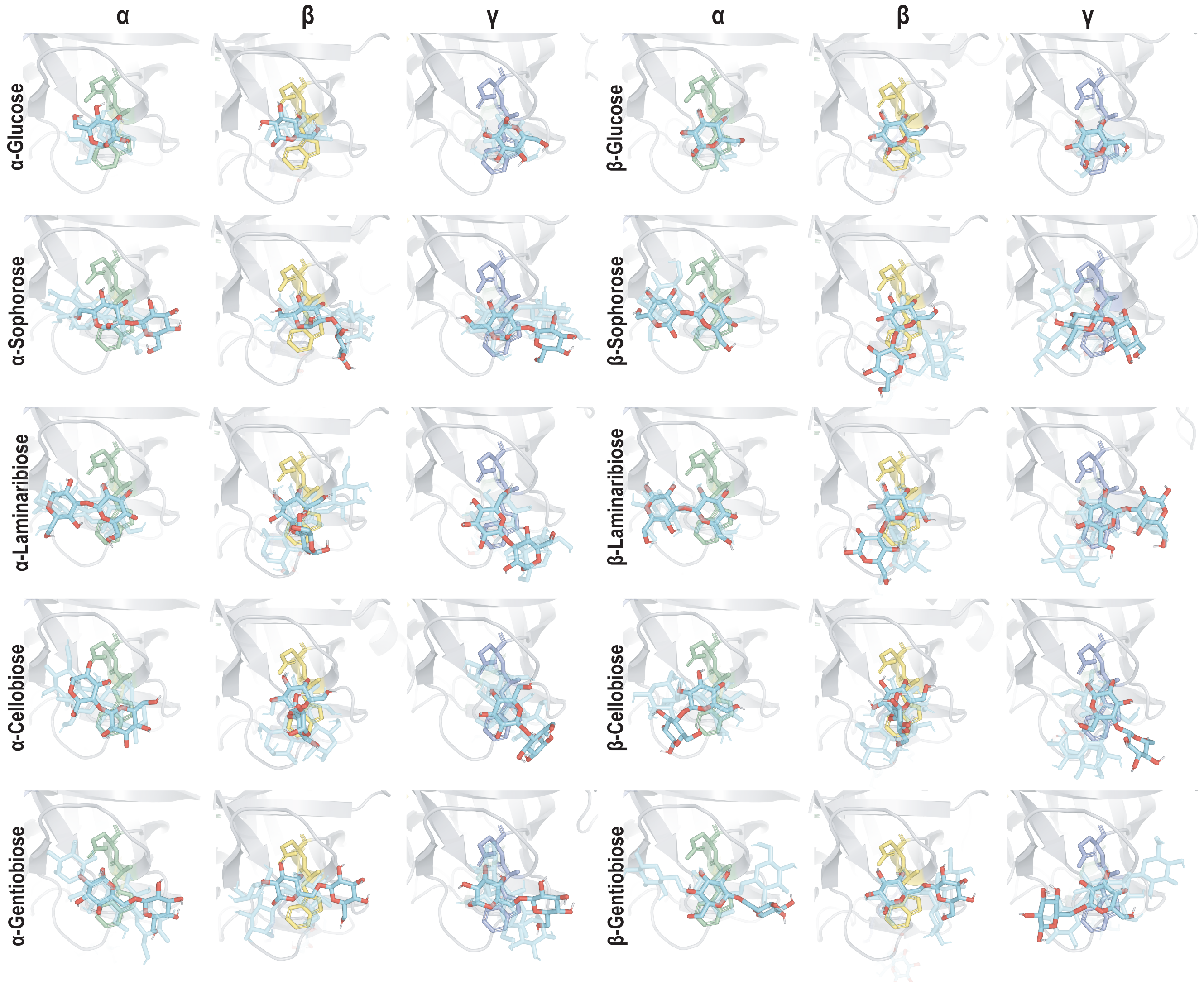


**Figure S11.** Predicted structures of *Cp*CBM92A α, β and γ binding sites in complex with glucose and disaccharide ligands generated using Boltz-1x and Gnina. For each ligand, the mode with the highest CNN score is depicted with red oxygen atoms, and the two next best-scoring modes are shown as transparent stick models to illustrate the variation between separate iterations of Gnina. The Trp-Glu binding sites are shown as green, yellow and blue sticks.

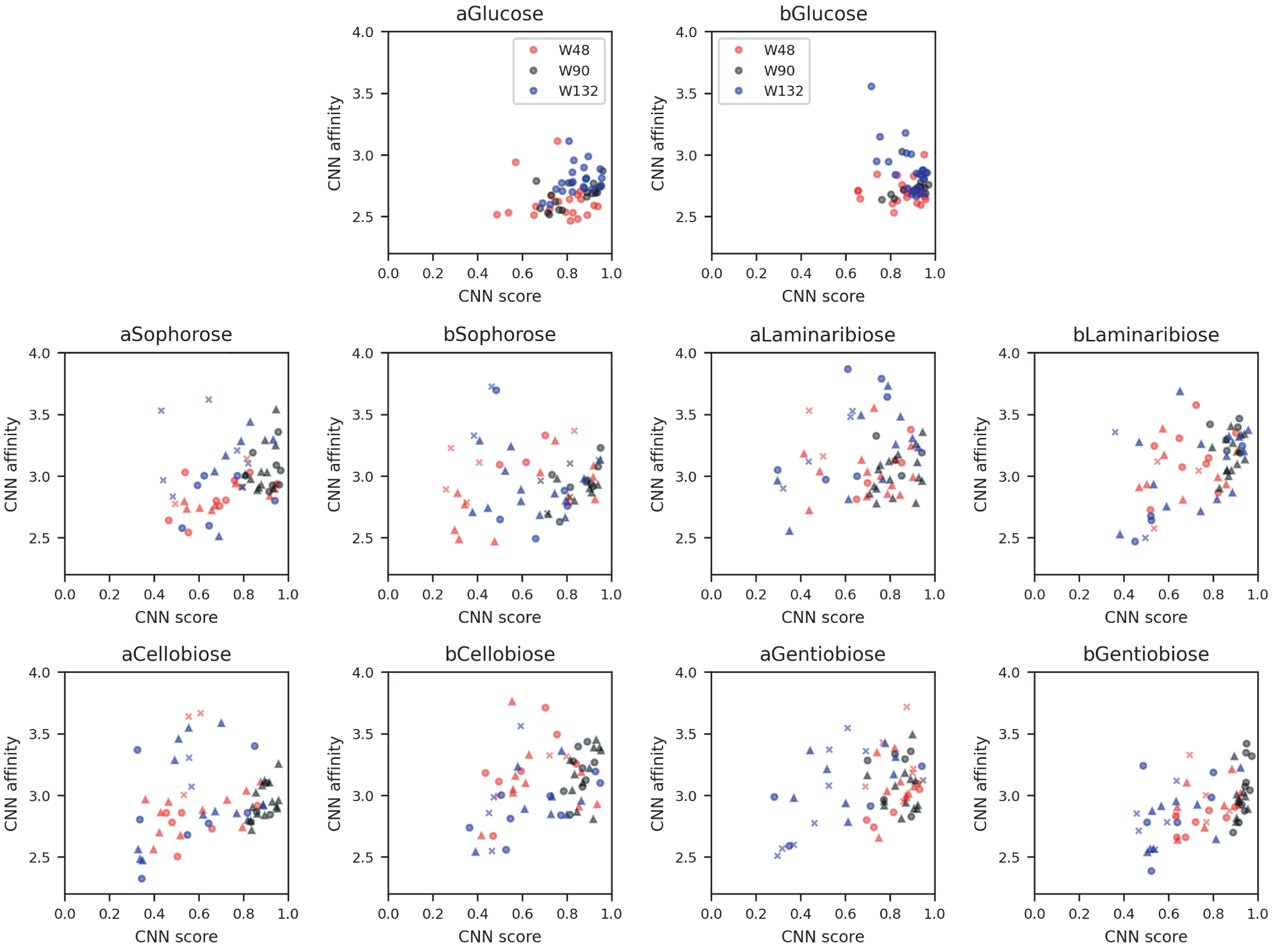


**Figure S12.** CNN scores and CNN affinities from dockings of glucose and disaccharide ligands in Gnina. The colors represent the α (W48, red), β (W90, black) and γ (W132, blue) binding sites. For disaccharides, the markers represent the glucosyl unit observed to interact directly with the Trp residue: circle = reducing end, triangle = non-reducing end, and cross = no glucosyl unit directly interacting with Trp.

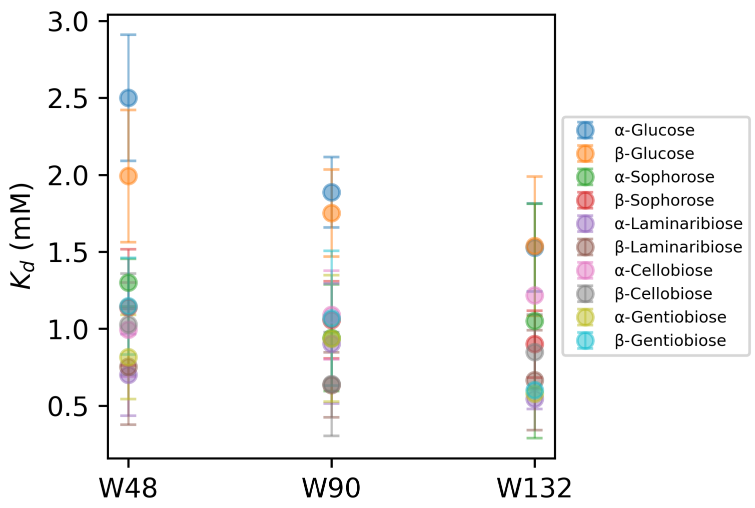


**Figure S13.** Predicted $K_D$ values for docked structures of glucose and disaccharide ligands at α, β and γ binding sites. The $K_D$ is calculated as the average of CNN affinities for structures output by Gnina with a CNN score >0.85. The error bars represent the standard deviation of the $K_D$.

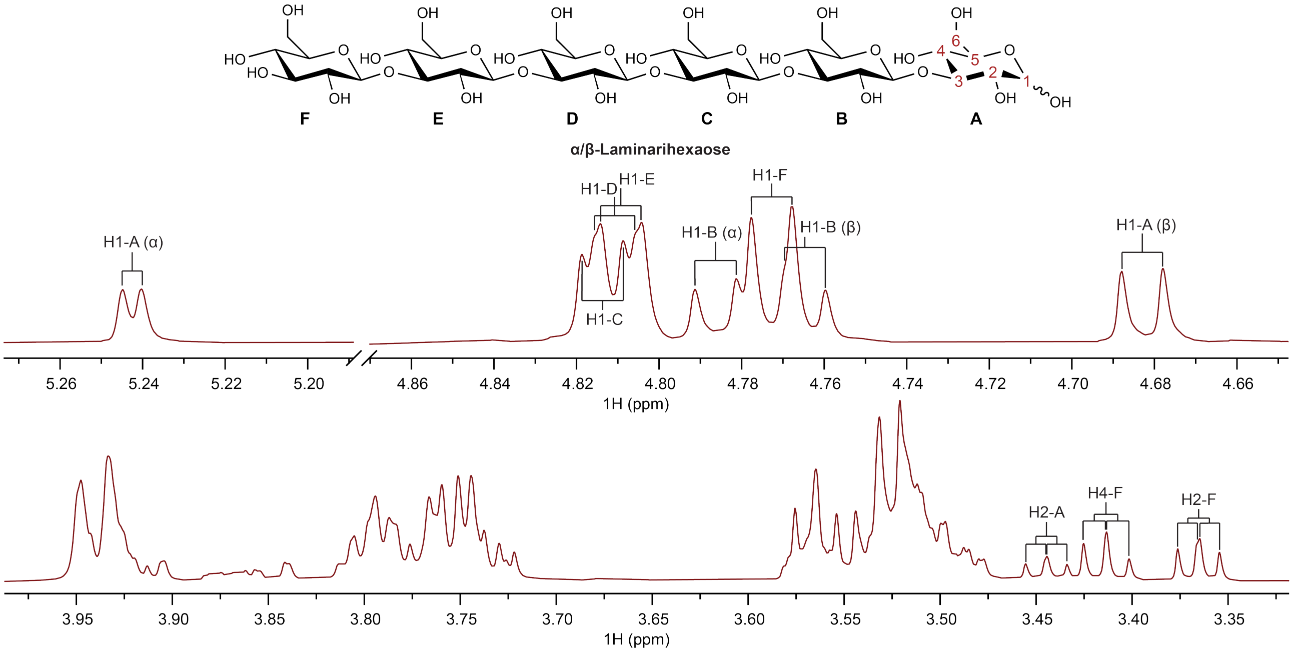


**Figure S14.** $^1$H NMR spectrum (800 MHz) of laminarihexaose acquired at 282 K. The spectrum is calibrated based on the TSP reference signal at 0 ppm.

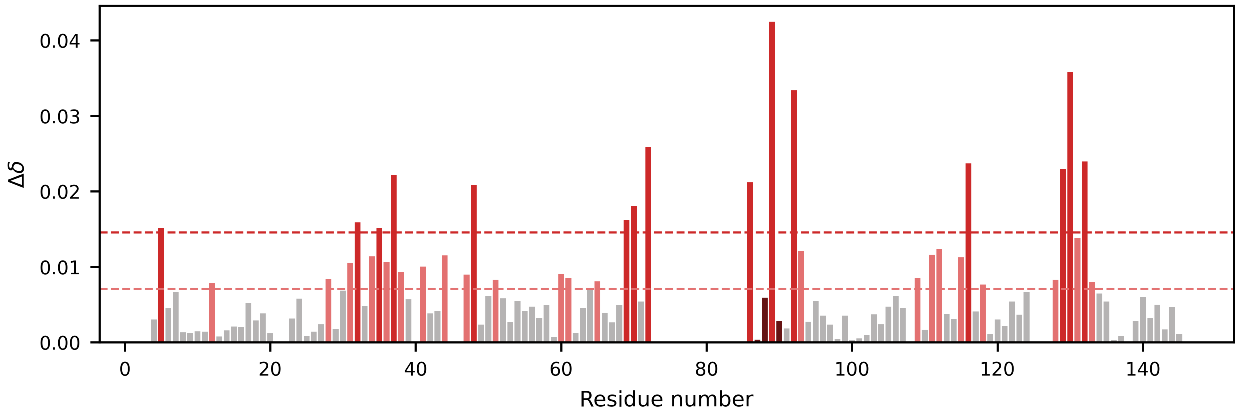


**Figure S15.** Chemical shift perturbations (CSP) of *Cp*CBM92A (80 µM) upon addition of laminarihexaose (800 µM). Horizontal lines and the corresponding color represent CSPs above the mean and one standard deviation above the mean CSP value. Lines in dark red represent residues with a high decrease in signal intensity upon additon of ligand.

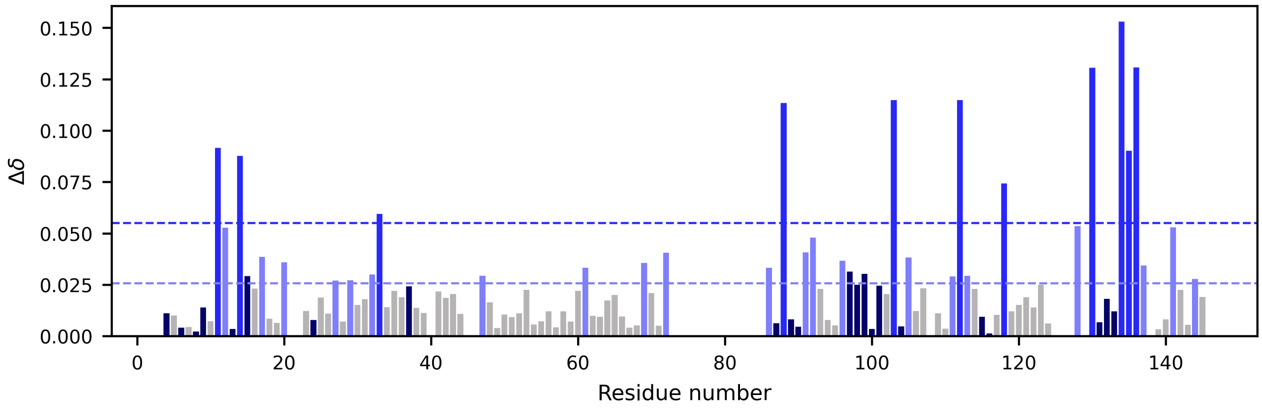


**Figure S16.** Chemical shift perturbations (CSP) of *Cp*CBM92A (80 µM) upon addition of gentiohexaose (800 µM). Horizontal lines and the corresponding color represent CSPs above the mean and one standard deviation above the mean CSP value. Lines in dark blue represent residues with a high decrease in signal intensity upon addition of ligand.

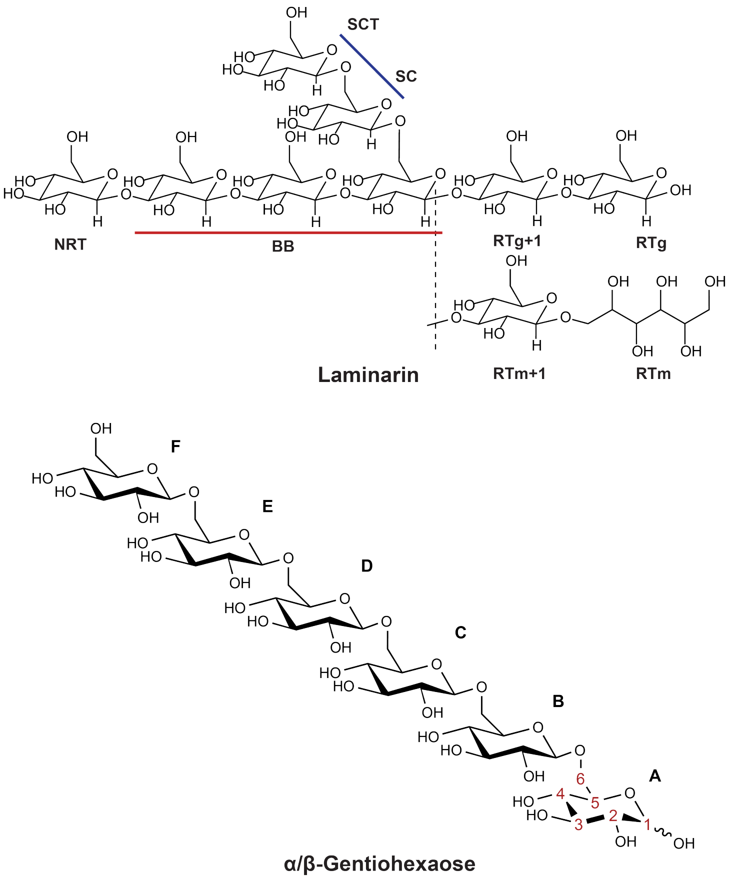


**Figure S17.** Model structure of laminarin (upper) showing the types of glucosyl units in the β-1,3-linked backbone (non-reducing end (NRT), main backbone (BB), G-series reducing end (RTg) and its neighbor (RTg+1), M-series reducing end (RTm) and its neighbor (RTm+1)) and the β-1,6-linked branches (sidechain non-terminal units (SC) and sidechain non-reducing ends (SCT)). The red and blue lines indicate where the chain may have varying lengths of glucosyl units in the backbone and side chain. Gentiohexaose structure (lower) with proton labels shown on the reducing end, and A-F labeling on glucosyl units.

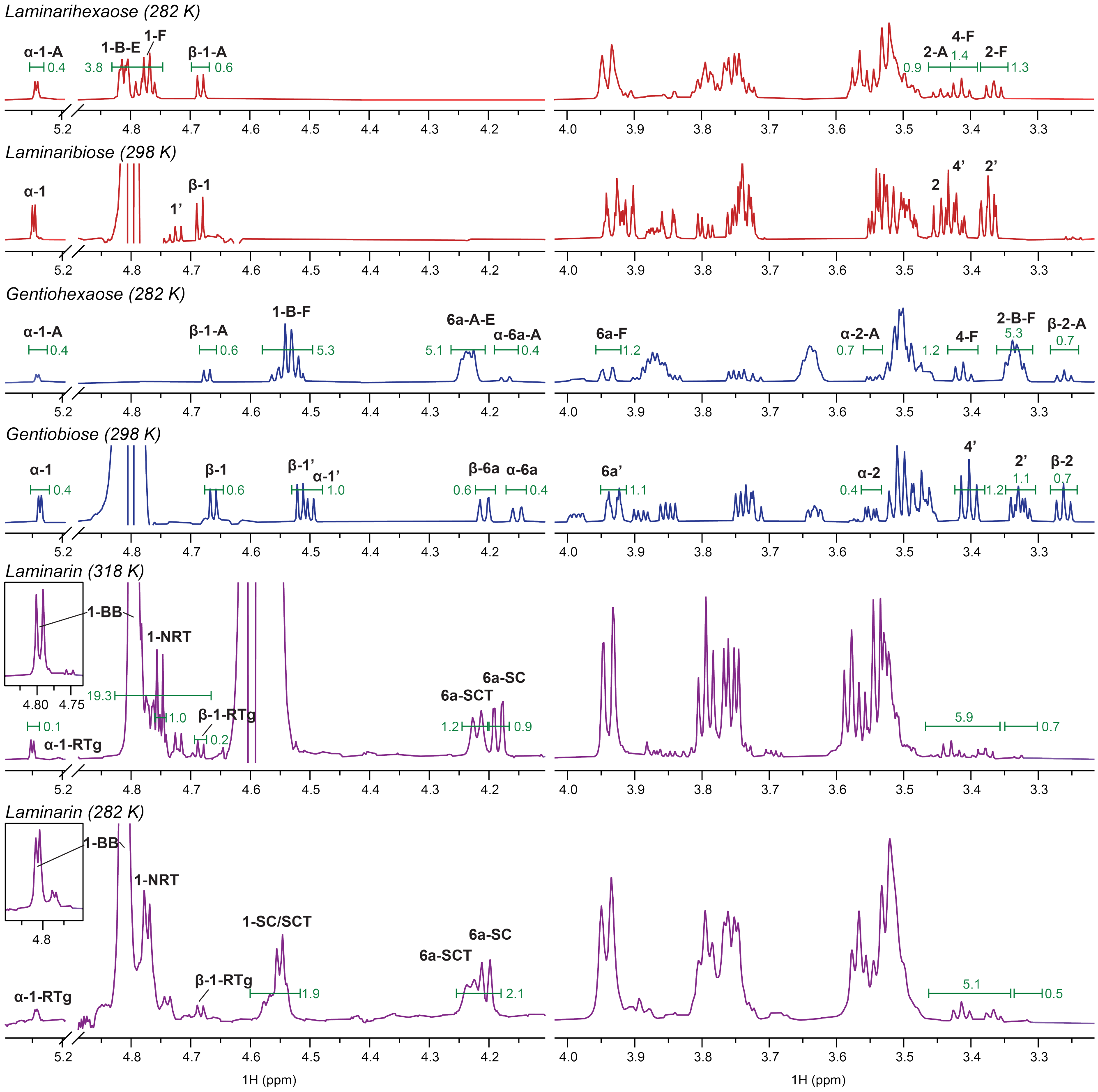


**Figure S18.** [1]H NMR spectra (800 MHz) laminarihexaose, laminaribiose, gentiohexaose, gentiobiose and laminarin acquired at temperatures specified in the figure. Integrals are shown with green intervals and numbers. Proton labels are based on annotations shown in ***Figure S17***. All spectra are calibrated based on the TSP reference signal at 0 ppm.

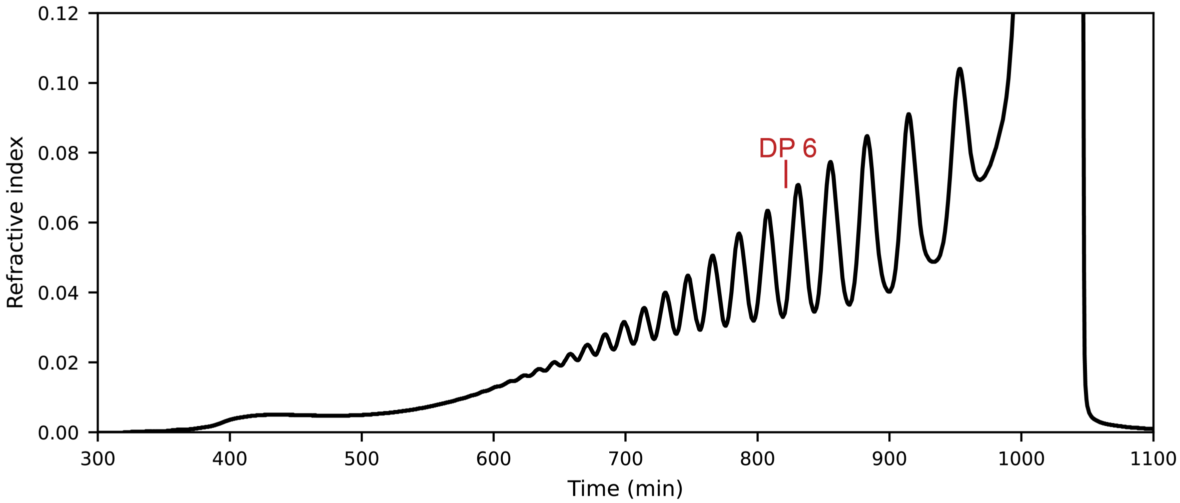


**Figure S19.** SEC chromatogram from gentiooligosaccharide separation on Superdex 30 column. The fraction containing gentiohexaose (DP 6) is indicated on the chromatogram.

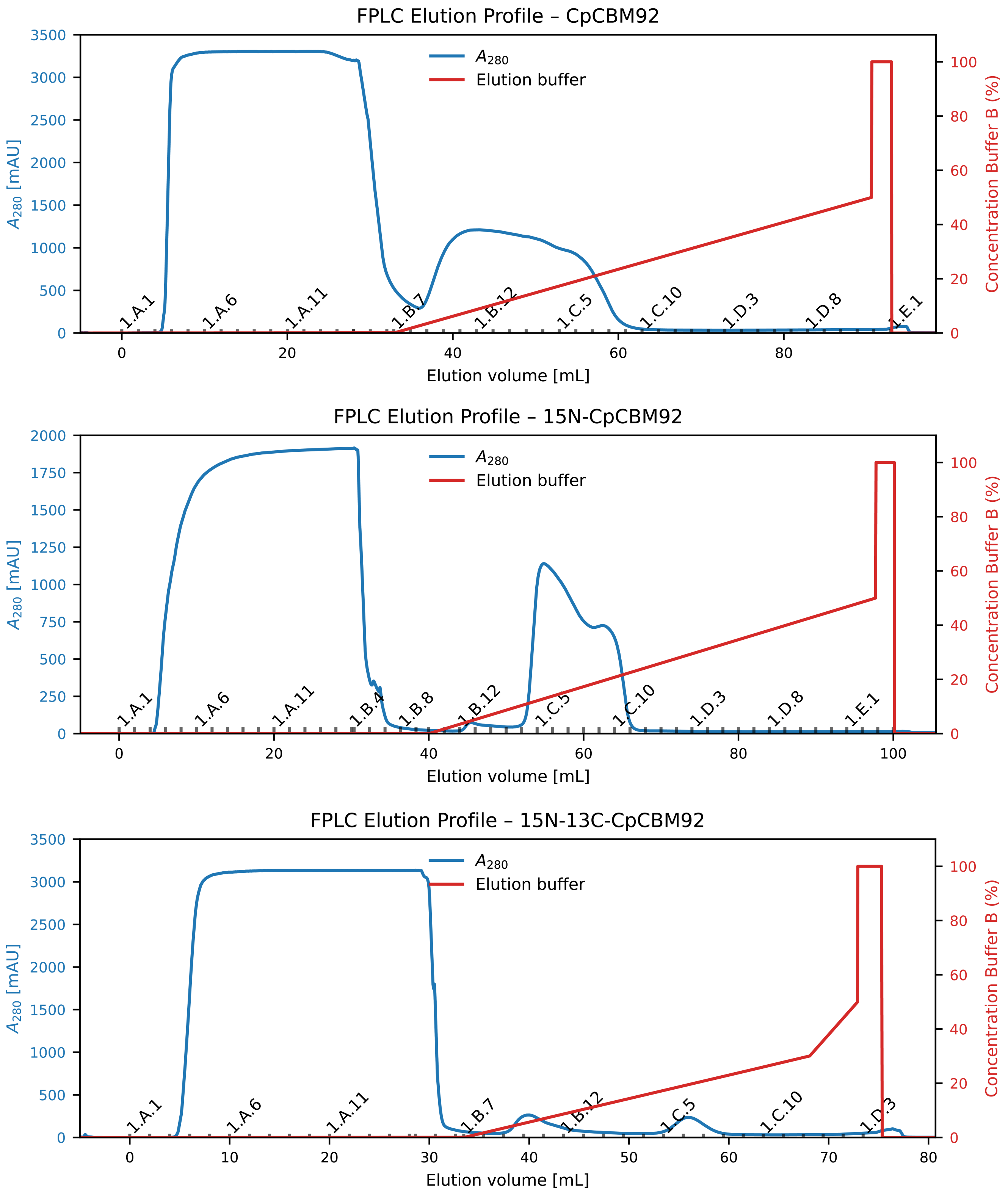


**Figure S20.** FPLC elution profiles from Ni affinity purification of non-labelled, $^{15}$N-labelled and $^{15}$N,$^{13}$C-labelled *Cp*CBM92A. Buffer B: 50 mM Tris-HCl (pH 8.0) supplemented with 300 mM NaCl and 400 mM imidazole.

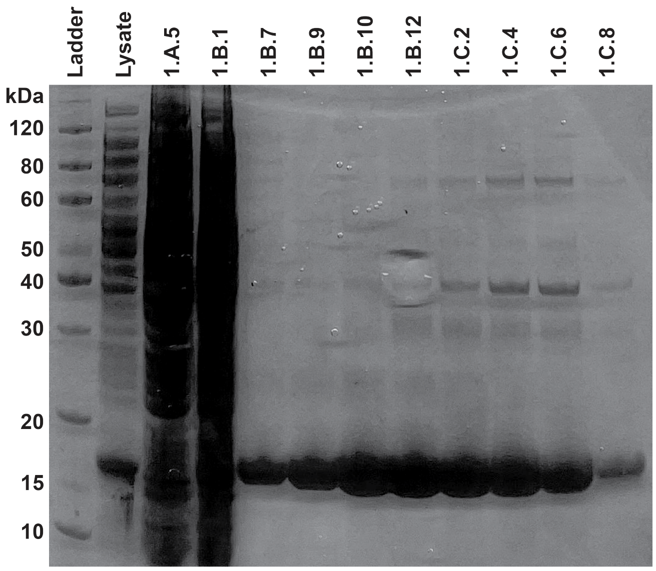


**Figure S21.** Picture of SDS-PAGE gel from purification of non-labelled *Cp*CBM92A (Mw = 16.2 kDa). The lanes are labelled with fraction numbers from FPLC as shown in ***Figure S20***.
